# Phylogenomic systematics of the spotted skunks (Carnivora, Mephitidae, *Spilogale*): Additional species diversity and Pleistocene climate change as a major driver of diversification

**DOI:** 10.1101/2020.10.23.353045

**Authors:** Molly M. McDonough, Adam W. Ferguson, Robert C. Dowler, Matthew E. Gompper, Jesús E. Maldonado

## Abstract

Four species of spotted skunks (Carnivora, Mephitidae, *Spilogale*) are currently recognized: *Spilogale angustifrons*, *S. gracilis*, *S. putorius*, and *S. pygmaea*. Understanding species boundaries within this group is critical for effective conservation given that regional populations or subspecies (e.g., *S. p. interrupta*) have experienced significant population declines. Further, there may be currently unrecognized diversity within this genus as some taxa (e.g., *S. angustifrons*) and geographic regions (e.g., Central America) never have been assessed using DNA sequence data. We analyzed species limits and diversification patterns in spotted skunks using multilocus nuclear (ultraconserved elements) and mitochondrial (whole mitogenomes and single gene analysis) data sets from broad geographic sampling representing all currently recognized species and subspecies. We found a high degree of genetic divergence among *Spilogale* that reflects seven distinct species and eight unique mitochondrial lineages. Initial divergence between *S. pygmaea* and all other *Spilogale* occurred 29 in the Early Pliocene (~ 5.0 million years ago) which was followed by subsequent diversification of the remaining *Spilogale* into an “eastern” and “western” lineage during the Early Pleistocene (~1.5 million years ago). These two lineages experienced temporally coincident patterns of diversification at ~0.66 and ~0.35 million years ago into two and ultimately three distinct evolutionary units, respectively. Diversification was confined almost entirely within the Pleistocene during a timeframe characterized by alternating glacial-interglacial cycles, with the origin of this diversity occurring in northeastern Mexico and the southwestern United States of America. Mitochondrial-nuclear discordance was recovered across three lineages in geographic regions consistent with secondary contact, including a distinct mitochondrial lineage confined to the Sonoran Desert. Our results have direct consequences for conservation of threatened populations, or species, as well as for our understanding of the evolution of delayed implantation in this enigmatic group of small carnivores.

## 1. Introduction

The taxonomy of well-known species is often taken for granted as being fully resolved, especially for less speciose and disproportionately studied groups such as mammals (Foley et al., 2016). In light of this, biologists studying such taxa tend to focus on genetic diversity below the species level, such as designation of Evolutionary Significant Units (Moritz, 1994) or genetically distinct groups or populations (Coates et al., 2018). This is especially true for North American carnivores, a well-studied group within which much work has been conducted on genetic distinctiveness at or below the species level (Akins et al., 2018; Balkenhol et al., 2020; Barton and Wisely, 2012; Nigenda-Morales et al., 2019; Wultsch et al., 2016). Although such work has highlighted taxonomic uncertainties, (e.g., Gopalakrishnan et al., 2018; Sinding et al., 2018), the overall alpha taxonomy of this group often remains unchanged and as such is readily perceived as more or less resolved.

For the many mammals threatened with extinction (Tilman et al., 2017), including carnivores and ungulates, changes of the underlying taxonomy can have direct consequences in terms of conservation efforts (Brownlow, 1996; Gippoliti et al., 2018; Thomson et al., 2018), with species often more valued from a conservation standpoint than are subspecies or local populations (Isaac et al., 2004; Mace, 2004). Examples where taxonomic uncertainty has the ability to threaten conservation efforts can be found in several threatened North American small carnivores including the pygmy raccoon *Procyon pygmaeus* and dwarf coati *Nasua nelson* (McFadden et al., 2008), island fox *Urocyon littoralis* (Hofman et al., 2015), and San Joaquin kit fox *Vulpes macrotis mutica* (Dragoo and Wayne, 2003). Although definitive boundaries for species recognition of some of these taxa appear debatable, especially from a genetic standpoint, biologists recognize the need to at minimum, treat these populations as distinct management units (Cuaron et al., 2009; Robertson et al., 2014). In the USA, for instance, the Endangered Species Act allows listing of species, subspecies, or Distinct Population Segments (DPSs) of vertebrates for special protections (Waples et al., 2018). Yet despite of the availability of multiple categories for protecting vertebrates as outlined in the ESA, assigning groups of animals to a particular category can have direct consequences on their conservation status (Gippoliti and Amori, 2007; Mace, 2004; O’Brien and Mayr, 1991) and can even influence taxonomic status or recognition (i.e., so-called ‘political species’; Karl and Bowen, 1999).

Accurate depictions of species diversity within a lineage also have consequences beyond conservation, including the understanding of their biogeography (Kreft and Jetz 2010; Escalante 2016), ecology (Isaac et al., 2004), and evolution (Upham et al., 2019). Species represent the fundamental unit in macroecology and inaccurate or out of date taxonomies can confound hypothesis testing (Isaac et al., 2004). For mammalian carnivores, macroecological studies on topics ranging from the relationship between geographic range size and body size (Diniz-Filho and Tôrres, 2002) to the evolution of delayed implantation (Thom et al., 2004) have often incorporated inaccurate information for one particular group of small carnivores: skunks.

Within the 12 currently recognized species of skunks in the family Mephitidae (Dragoo, 2009), molecular-based studies have mostly focused on unravelling their inter-generic relationships (Dragoo et al., 1993) or focused specifically on the systematics of hog-nosed skunks in the genus *Conepatus* (Dragoo et al., 2003; Schiaffini et al., 2013). However, to date, the evolutionary history and phylogenetic relationships of the different species of the genus *Spilogale* Gray 1865 in particular have been poorly studied. As a result, the genus has been plagued by taxonomic uncertainty since it was first described (Table 1). These mephitids are distributed north to south from British Columbia to Costa Rica and east to west from West Virginia to California, and inhabit a wide array of biogeographic regions with complex geological and climatic histories. The four currently-recognized species (*S. angustifrons*, *S. gracilis*, *S. putorius*, *S. pygmaea*; Dragoo, 2009) inhabit the following North American biogeographic areas that have been well-characterized for mammals: Californian-Rocky Mountain subregion (Escalante et al., 2013), northwestern North America (Shafer et al., 2010), the warm deserts (Riddle and Hafner, 2006), the Great Basin (Riddle et al., 2014), the Mexican Transition Zone (Morrone, 2010; Morrone, 2015), the Alleghanian subregion (Escalante et al., 2013), and Central America (Gutiérrez-García and Vázquez-Domínguez, 2017).

**Table 1.**
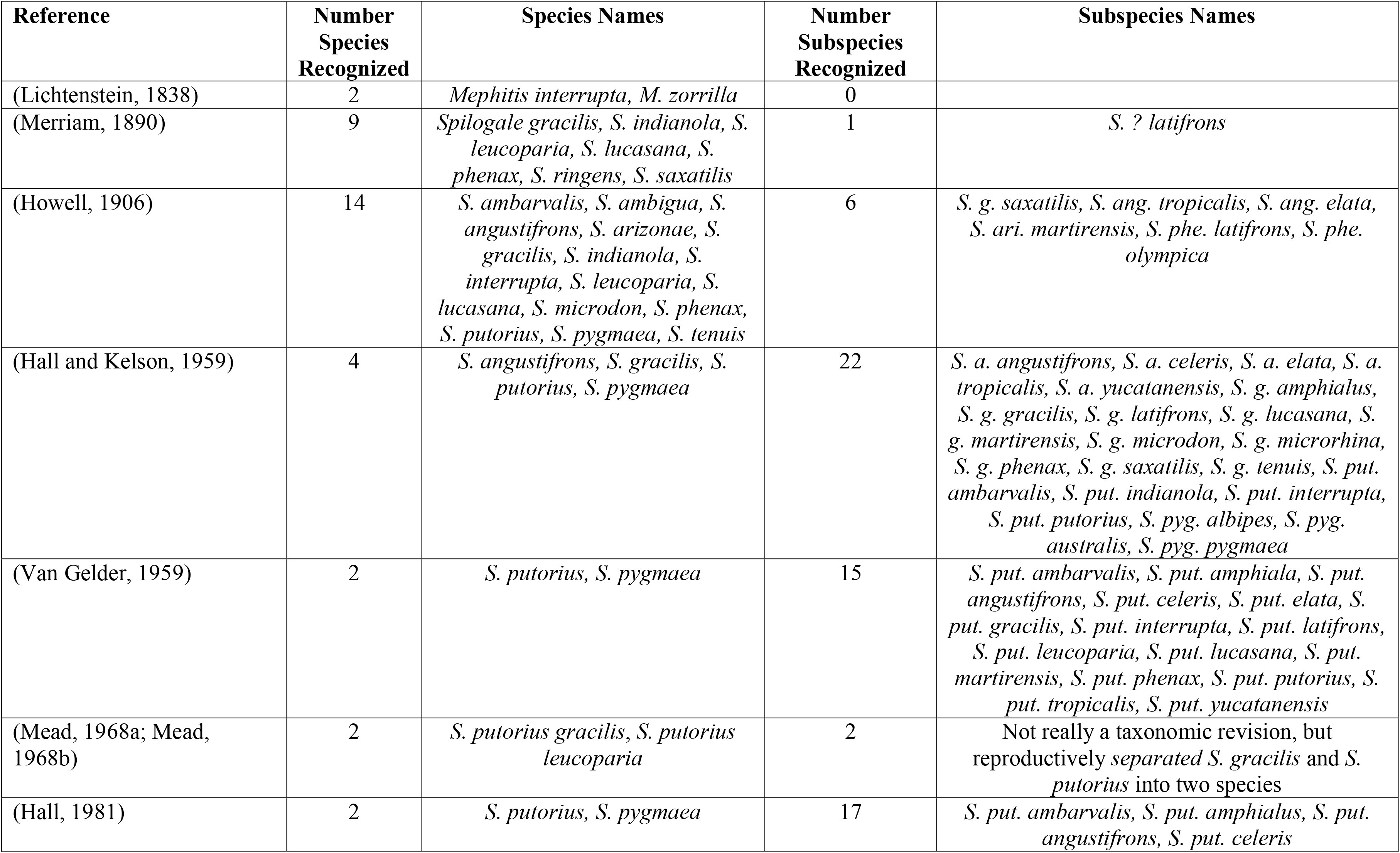
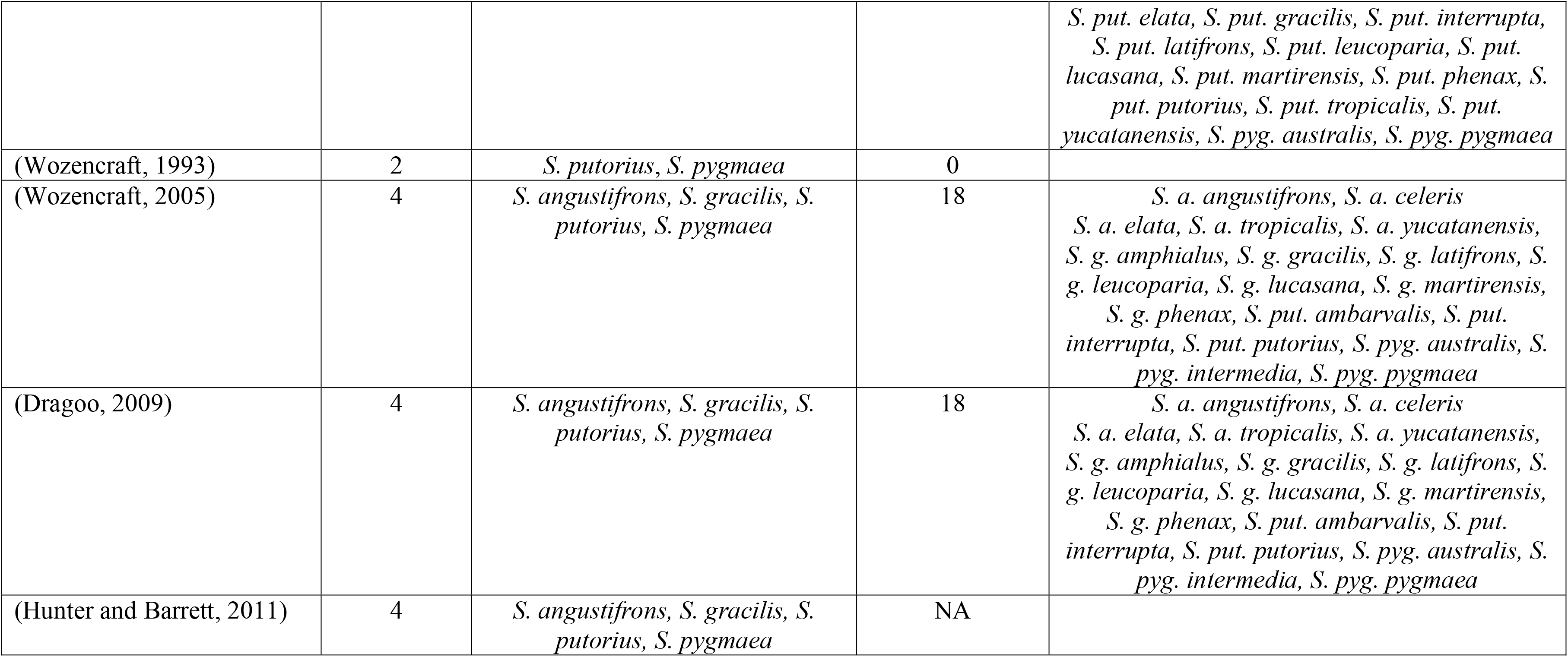
Summaries of major taxonomic revisions and mammalian taxonomic references highlighting the numbers of species and subspecies within the genus *Spilogale* (Mammalia, Carnivora, Mephitidae).

| Reference | Number<br>Species<br>Recognized | Species Names | Number<br>Subspecies<br>Recognized | Subspecies Names |
| --- | --- | --- | --- | --- |
| (Lichtenstein, 1838) | 2 | *Mephitis interrupta*, *M. zorrilla* | 0 |  |
| (Merriam, 1890) | 9 | *Spilogale gracilis*, *S. indianola*, *S. leucoparia*, *S. lucasana*, *S. phenax*, *S. ringens*, *S. saxatilis* | 1 | *S. ? latifrons* |
| (Howell, 1906) | 14 | *S. ambarvalis*, *S. ambigua*, *S. angustifrons*, *S. arizonae*, *S. gracilis*, *S. indianola*, *S. interrupta*, *S. leucoparia*, *S. lucasana*, *S. microdon*, *S. phenax*, *S. putorius*, *S. pygmaea*, *S. tenuis* | 6 | *S. g. saxatilis*, *S. ang. tropicalis*, *S. ang. elata*, *S. ari. martirensis*, *S. phe. latifrons*, *S. phe. olympica* |
| (Hall and Kelson, 1959) | 4 | *S. angustifrons*, *S. gracilis*, *S. putorius*, *S. pygmaea* | 22 | *S. a. angustifrons*, *S. a. celeris*, *S. a. elata*, *S. a. tropicalis*, *S. a. yucatanensis*, *S. g. amphialus*, *S. g. gracilis*, *S. g. latifrons*, *S. g. lucasana*, *S. g. martirensis*, *S. g. microdon*, *S. g. microrhina*, *S. g. phenax*, *S. g. saxatilis*, *S. g. tenuis*, *S. put. ambarvalis*, *S. put. indianola*, *S. put. interrupta*, *S. put. putorius*, *S. pyg. albipes*, *S. pyg. australis*, *S. pyg. pygmaea* |
| (Van Gelder, 1959) | 2 | *S. putorius*, *S. pygmaea* | 15 | *S. put. ambarvalis*, *S. put. amphiala*, *S. put. angustifrons*, *S. put. celeris*, *S. put. elata*, *S. put. gracilis*, *S. put. interrupta*, *S. put. latifrons*, *S. put. leucoparia*, *S. put. lucasana*, *S. put. martirensis*, *S. put. phenax*, *S. put. putorius*, *S. put. tropicalis*, *S. put. yucatanensis* |
| (Mead, 1968a; Mead, 1968b) | 2 | *S. putorius gracilis*, *S. putorius leucoparia* | 2 | Not really a taxonomic revision, but reproductively separated *S. gracilis* and *S. putorius* into two species |
| (Hall, 1981) | 2 | *S. putorius*, *S. pygmaea* | 17 | *S. put. ambarvalis*, *S. put. amphialus*, *S. put. angustifrons*, *S. put. celeris* |
|  |  |  |  | *S. put. elata, S. put. gracilis, S. put. interrupta, S. put. latifrons, S. put. leucoparia, S. put. lucasana, S. put. martirensis, S. put. phenax, S. put. putorius, S. put. tropicalis, S. put. yucatanensis, S. pyg. australis, S. pyg. pygmaea* |
| (Wozencraft, 1993) | 2 | *S. putorius, S. pygmaea* | 0 |  |
| (Wozencraft, 2005) | 4 | *S. angustifrons, S. gracilis, S. putorius, S. pygmaea* | 18 | *S. a. angustifrons, S. a. celeris S. a. elata, S. a. tropicalis, S. a. yucatanensis, S. g. amphialus, S. g. gracilis, S. g. latifrons, S. g. leucoparia, S. g. lucasana, S. g. martirensis, S. g. phenax, S. put. ambarvalis, S. put. interrupta, S. put. putorius, S. pyg. australis, S. pyg. intermedia, S. pyg. pygmaea* |
| (Dragoo, 2009) | 4 | *S. angustifrons, S. gracilis, S. putorius, S. pygmaea* | 18 | *S. a. angustifrons, S. a. celeris S. a. elata, S. a. tropicalis, S. a. yucatanensis, S. g. amphialus, S. g. gracilis, S. g. latifrons, S. g. leucoparia, S. g. lucasana, S. g. martirensis, S. g. phenax, S. put. ambarvalis, S. put. interrupta, S. put. putorius, S. pyg. australis, S. pyg. intermedia, S. pyg. pygmaea* |
| (Hunter and Barrett, 2011) | 4 | *S. angustifrons, S. gracilis, S. putorius, S. pygmaea* | NA |  |

The taxonomy of *Spilogale* has been rife with change across the 19^th^ and 20^th^ centuries, with as few as two (Lichtenstein, 1838) to as many as 14 (Howell, 1906) species recognized at any given time (Table 1). Major taxonomic revisions that attempted to determine the number of spotted skunk species, mostly based on morphological characters, have used a combination of craniodental features and pelage patterns (Howell, 1906; Van Gelder, 1959). However, this system of classification has proved problematic due to the conserved external morphology of spotted skunks and intergradation of characters across proposed taxonomic boundaries (Van Gelder, 1959). Species limits within spotted skunks were further confounded by studies that revealed differences in their reproductive biology including the presence or absence of delayed implantation for particular subspecies (Mead, 1968a; Mead, 1968b). Within the genus *Spilogale*, most molecular studies to date have focused on intra-specific patterns of genetic diversity including the recognition of the genetic distinctness of insular populations (Floyd et al., 2011; Jones et al., 2013), influences of Pleistocene climate change on phylogeographic subdivisions (Ferguson et al., 2017), and subspecies boundaries (Shaffer et al., 2018). Recent focus on populations of the plains spotted skunk subspecies, *S. putorius interrupta*, are particularly relevant given conservation concerns and a recent petition to have this subspecies listed as endangered under the Endangered Species Act (USFWS, 2012) and documented range-wide decline across much of the Great Plains (Gompper, 2017; Gompper and Hackett, 2005). Thus, understanding the taxonomy of *S. putorius* is especially important in light of this proposed conservation action and particularly relevant considering the strong genetic subdivision documented among the three recognized *S. putorius* subspecies (Shaffer et al., 2018). Understanding the evolutionary uniqueness of *S. putorius interrupta* in particular, and all spotted skunks in general, requires a detailed molecular study of all *Spilogale* species with samples from across their distribution. Such an effort would also shed light on the taxonomic status of one of the most enigmatic species of the group: *S. angustifrons*.

The southern spotted skunk, *S. angustifrons* was first recognized as a full species by Howell (1902) and periodically appeared or was removed from the taxonomically valid list of spotted skunks over the next century (Table 1). Its recognition as a valid species was based solely on morphological characters (body and skull size, spotting pattern) until the late 1990s when two individuals from El Salvador were karyotyped, providing the only genetic data on this species to date (Owen et al., 1996). Since then, most recent treatises on mammal taxonomy have recognized *S. angustifrons* as a distinct species (Dragoo, 2009; Wozencraft, 2005), which is purported to range from central Mexico south to Costa Rica (Reid, 2009). In the absence of modern molecular data, the validity of *S. angustifrons* as well as other questions important to conserving populations of spotted skunks cannot be fully realized.

Here we present the first complete molecular phylogeny of the genus *Spilogale* using genomic data from ultraconserved elements (UCEs) and complete mitogenomes from a combination of modern and historical museum specimens representing all four currently recognized species and their associated subspecies. Our main objective was to test hypotheses regarding species limits within *Spilogale* with a particular emphasis on the validity of *S. angustifrons* and genetic uniqueness of *S. putorius interrupta*. To test these hypotheses we: (1) generated concatenated and species-tree phylogenies for all species of *Spilogale* from across their entire range; (2) assessed the influence of historical processes associated with geologic and climatic processes on genetic structure using time-calibrated phylogenies; and (3) inferred historical patterns of range shifts associated with identified species or genetically distinct populations. Our results have important implications for the taxonomy and conservation of spotted skunks across North America and provide strong evidence for the presence of seven species. Proposed taxonomic recommendations provided herein, in conjunction with reassessments of the distributions and conservation status of these species, will facilitate more effective management and provide additional insight into the important role that the evolution of different reproductive strategies such as the presence or absence of delayed implantation had on the diversification patterns observed in this group.

## 2. Materials and Methods

### 2.1. Sampling and DNA extraction

Using a combination of recently collected tissues (i.e., modern samples) and destructively sampled material from museum voucher specimens (i.e., historic samples) we assembled a collection of 203 spotted skunk samples (n = 177 modern and n = 26 historic) from throughout most of the range of the genus (Fig. 1, Fig. 2, Supplementary Table S1). Our samples included representatives of the four currently recognized species of spotted skunks as well as all currently recognized subspecies with the exception of *S. pygmaea intermedia* (Dragoo 2009, Supplementary Table S1): the southern spotted skunk *S. angustifrons* (*S. a. angustifrons*, *S. a. elata*, *S. a. celeris*, *S. a. tropicalis*, and *S. a. yucatanensis*), the western spotted skunk *S. gracilis* (*S. g. amphialus*, *S. g. gracilis*, *S. g. latifrons*, *S. g. leucoparia*, *S. g. lucasana*, *S. g martirensis*, *S. g. phenax*), the eastern spotted skunk *S putorius* (*S. put. ambarvalis*, *S. put. interrupta*, *S. put. putorius*), and the pygmy spotted skunk *S. pygmaea* (*S. pyg. australis*, *S. pyg. intermedia*, and *S. pyg. pygmaea*). For outgroup comparison, we obtained samples from two North American hog-nosed skunks (*Conepatus leuconotus*), one striped hog-nosed skunk (*Conepatus semistriatus*), two striped skunks (*Mephitis mephitis*), one hooded skunk (*Mephitis macroura*), and one Javan stink badger (*Mydaus javanensis*). Samples were sourced from 28 natural history museums and a diversity of researchers from across North America (see Acknowledgements). For specimens collected as part of this study, all animals were taken according to the guidelines of the American Society of Mammalogists for use of wild mammals in research (Sikes et al., 2016) and adhered to Institutional Animal Care and Use Committee (IACUC) standards from the respective institutions.

**Figure 1.**
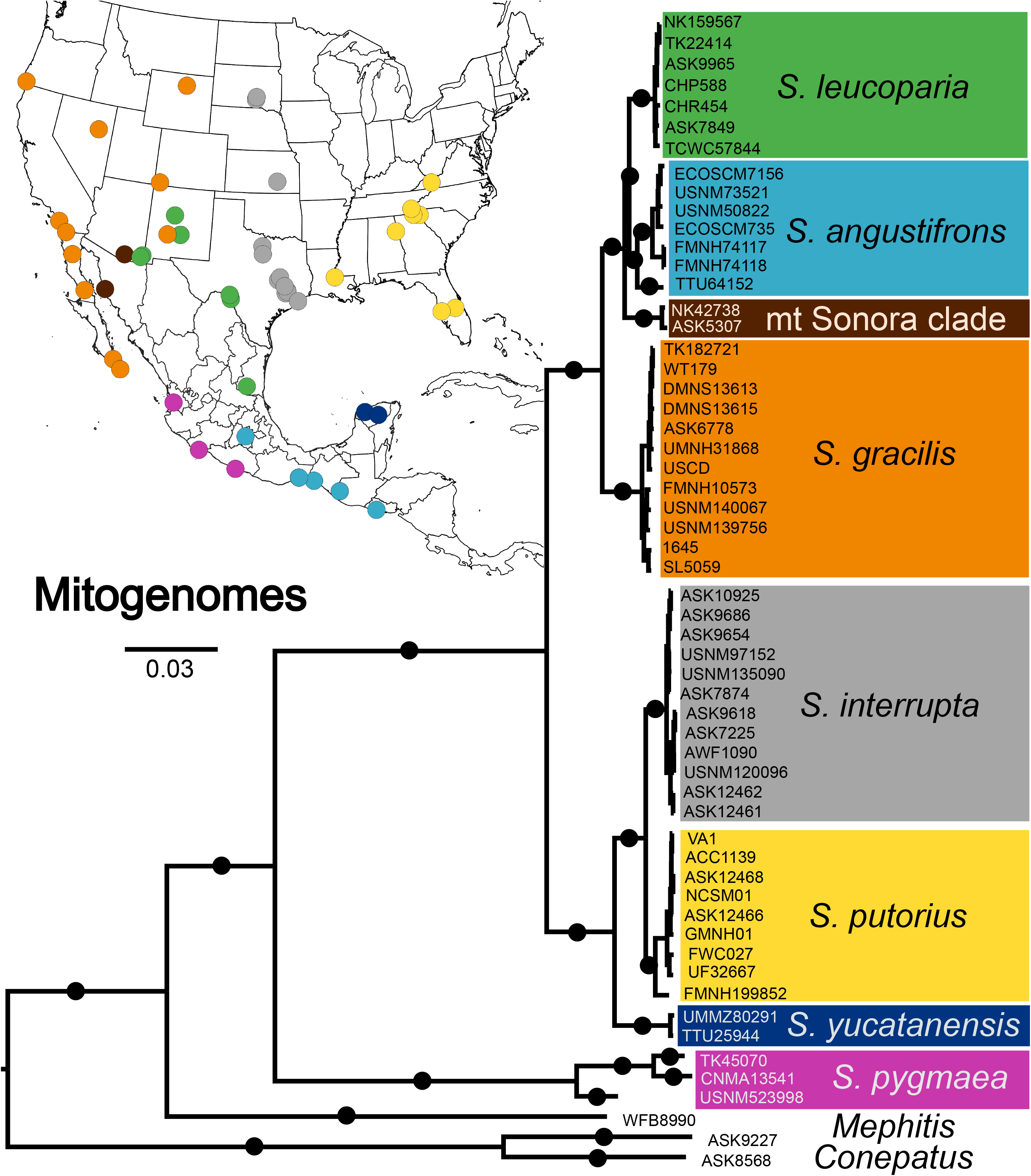
Bayesian phylogeny of *Spilogale* species based on analysis of mitochondrial genomes. Black circles on the branches leading to the major clades indicated Bayesian posterior probabilities = 1 and bootstrap values > 90. Map inset includes sampling localities for mitogenomes with colored dots corresponding to major clades.

**Figure 2.**
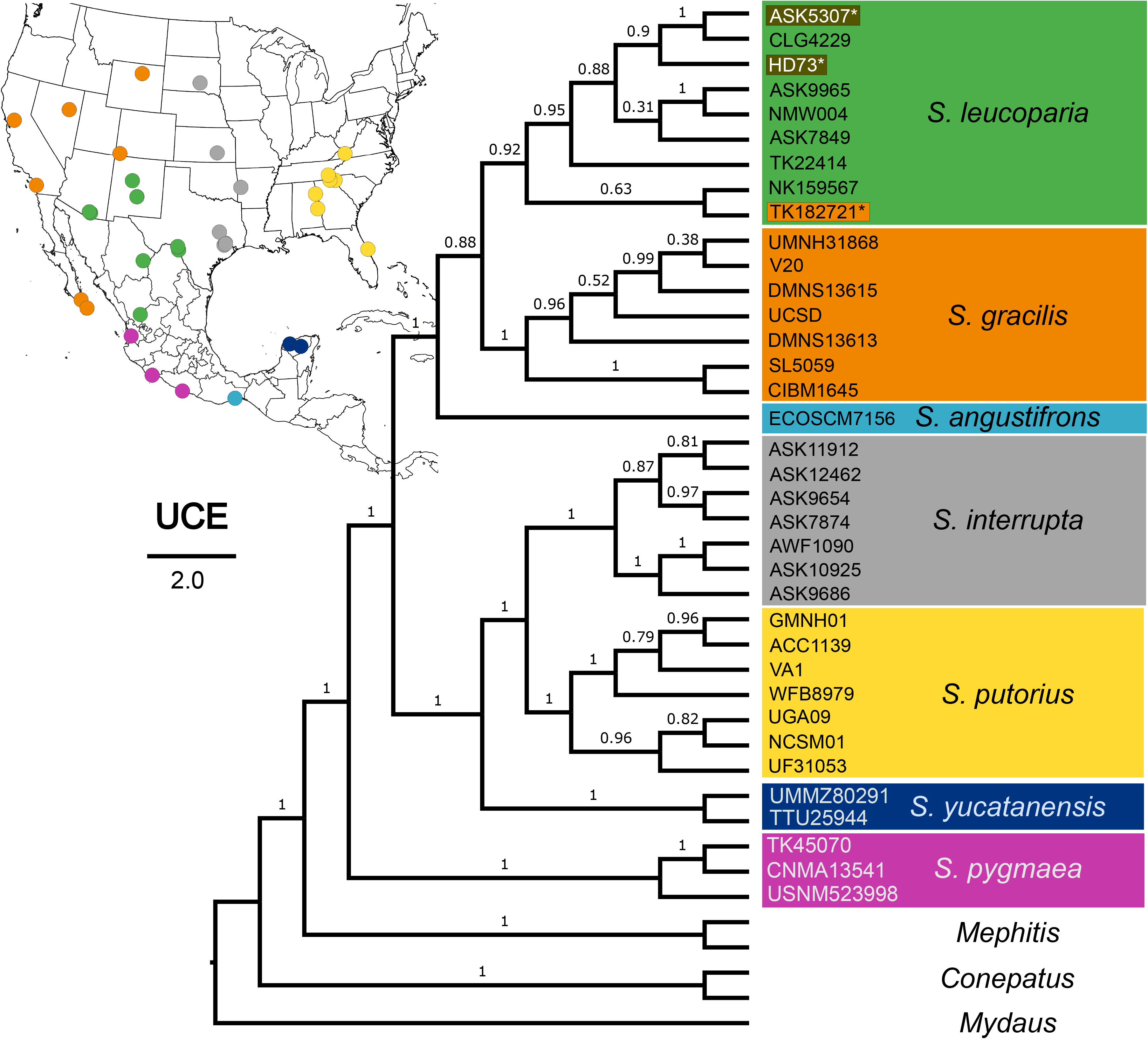
Cladogram of *Spilogale* species based on analysis of 3,896 UCE loci (60% matrix). Quartet values from species tree are displayed along the branches. Map inset includes sampling localities for UCE.

DNA was extracted from frozen or ethanol preserved tissue with a DNeasy blood and tissue kit (Qiagen) following manufacturers protocols. DNA was quantified using a Qubit dsDNA High Sensitivity Assay kit and samples were diluted to 200 ng in 50 μL. We sheared the samples to a targeted fragment size ~150-350 bp with Covaris microTUBES.

DNA isolation from historical samples (source material mainly from claws of dried museum skins) followed the digestion and phenol-chloroform extraction protocols outlined in McDonough et al. (2018). For the historical samples, DNA was not sheared and irrespective of the starting concentration, 35 μL of DNA was aliquoted and concentrated using a 3X bead clean-up with speedbead recipe following Rohland and Reich (2012) and resuspended in 14 μL of DNAse free water.

### 2.2. Library preparation and targeted enrichment

KAPA LTP (Kapa Biosystems) library preparation for Illumina platforms followed the manufacturer’s protocol but were modified to use one-quarter of the reagents per reaction. Amplifications were performed using 15 cycles with Nextera-style dual indexing primers (Kircher et al., 2011) and KAPA HiFi Hotstart ReadyMix (Kapa Biosystems). Library concentrations were measured with Qubit dsDNA High Sensitivity Assay kit.

For a subset of individuals with DNA sourced from well-preserved tissue, the entire mitochondrial genome was amplified using two sets of universal primers: 1) tRNA Leuc1 and *16S* rRNA1, and 2) tRNA Leuc2 and *16S* rRNA2 (Sasaki et al., 2005) and libraries were prepared using NexteraXT following Ferguson et al. (2019).

Libraries were enriched for UCEs with the 5K v1 Tetrapods myBaits kit (Faircloth et al., 2012) produced by Arbor Biosystems (Ann Arbor, MI) using 5,472 baits (120 mer) that target 5,060 UCEs. We followed the manufacturer’s protocols, including longer hybridization time for historical samples (18-24 hours). Prior to enrichment, samples were pooled in equimolar ratios with up to 8 libraries for modern and up to 4 libraries for historical samples before enrichment.

For high-quality DNA samples, entire mitochondrial genomes are often recovered as a byproduct of UCE enrichment; however, full recovery from degraded DNA samples such as those from historical museum specimens is often not possible. Therefore, the libraries from historical samples were separately enriched for mitochondrial genomes using a custom bait set designed from mephitid mitogenome sequences available on GenBank (*Spilogale putorius*, AM711898) to produce a 20,000 probe myBaits kit and following Hawkins et al. (2016). Post-capture amplification was performed using Illumina primers for 15 cycles and final products were quantified using Qubit and pooled in equal nanomolar concentrations. Sequencing was performed by Novogene Corp. Inc. on a HiSeq4000 Illumina system.

### 2.3. Mitogenome processing and phylogenetic methods

After demultiplexing, we combined paired-end reads using FLASH2 v. 2.2.00 (Magoč and Salzberg, 2011) and trimmed adapter sequences using TrimGalore v. 0.4.3 (Krueger, 2015). Poor quality sequences (scores below 20) and exact PCR replicates were removed using prinseq-lite v.0.20.4 (Schmieder and Edwards, 2011). Reads were mapped to a previously sequenced *Spilogale* mitochondrial genome (GenBank number AM711898) using the Burrows-Wheeler algorithm in BWA v.7.10 (Li and Durbin, 2009) using the “bwa mem” command. We imported mapped reads into Geneious Prime v. 2020.2.2 (Biomatters LTD) and examined the assemblies by eye. The reference genome (above) was used to transfer genome annotations. To rule out the presence of nuclear copies of mitochondrial genes (NUMTs), we translated all protein-coding genes to check for frame shifts or stop codons. Mitochondrial genomes are available on GenBank under the accession numbers: XXXX-XXXX.

We created alignments using the entire mitogenome with Muscle v. 3.8.425 (Edgar, 2004) using 8 iterations. We used the concatenated alignment (without partitions) for estimating the phylogenetic relationships within *Spilogale* using RAxML v. 8 (Stamatakis, 2014). The tree with the best likelihood score was selected from 1,000 trees generated using the GTRGAMMA nucleotide substitution model. Bootstrap values were obtained using 1,000 replicates under the GTRGAMMA substitution model. Finally, -f b command was used to create the bipartitions file that assigned support values to branches.

Bayesian inference (BI) was conducted on a partitioned dataset using MrBayes v. 3.2.6 (Ronquist and Huelsenbeck, 2003). The best partition scheme was estimated using the greedy search algorithm in PartionFinder2 version. (Lanfear et al., 2016). Our search was limited to those models available in MrBayes and using the corrected Akaike information criterion (AICc). The data block was defined using 39 partitions: 13 protein-coding genes, 22 tRNAs, two rRNAs, the origin of replication, and the control region and the codons were partitioned for each of the protein coding genes. The BI was performed using four Markov-chain Monte Carlo (MCMC) runs with three heated and one cold chain. Two independent runs were conducted with 50 million generations, sampling trees and parameters every 5000 generations, and a burn-in of 10% of the sampled trees. Convergence of MCMC runs were verified using Tracer v. 1.7 (Rambaut et al., 2018). Both BI and Maximum likelihood (ML) estimates were implemented on the CIPRES science gateway (www.phylo.org). The final trees and support values were visualized using FigTree v.1.4.4 (http://tree.bio.ed.ac.uk/software/figtree/).

### 2.4. Processing of UCEs and phylogenetic analyses

We followed the Phyluce v. 1.6 pipeline (Faircloth, 2015) https://github.com/faircloth-lab/phyluce to process the raw reads. We used Illumiprocessor v. 2.0.7 (Faircloth, 2013) and Trimmomatic v. 0.32 (Bolger et al., 2014) to trim adapters and low-quality regions. We used Trinity v. 2.0.6 (Grabherr et al., 2011) to assemble the contigs. Next, contigs were matched to the 5,060 tetrapod UCE probes. We used the Phyluce script phyluce_align_seqcap_align for edge-trimming and we aligned UCE loci using MAFFT 7 (Katoh and Standley, 2013). Finally, we tested several alignments that included various degrees of missing data: 50% matrix for which half of the taxa were present for each UCE locus, 60% matrix (40% of taxa missing), and 70% matrix (30% of taxa missing).

We conducted a ML analysis on the concatenated alignments using RAxML v. 8 (Stamatakis, 2014) on the CIPRES Science Gateway v. 3.3 (https://www.phylo.org) with 500 rapid nonparametric bootstrap replicates using the GTRCAT approximation.

For the species tree analysis, we inferred individual gene trees using ParGenes v. 1.0.1 (Morel et al., 2018) with model selection using ModelTest-NG (Darriba et al., 2019) and tree inference using RAxML-NG (Kozlov et al., 2019). Coalescent-based species tree analysis was performed using ASTRAL-III v. 5.6.3 (Zhang et al., 2018). The UCE alignments and analyses files are available from the Dryad Digital Repository (http://datadryad.org/XXXX)

### 2.5. Mitochondrial DNA analyses

#### 2.5.1 Divergence time estimates

Divergence time estimates were implemented in BEAST v. 2.6.1 (Bouckaert et al., 2019) using a concatenated alignment of the 13 protein coding genes in the mitochondrial genome. The alignment included a subset of samples representing two individuals from each of the clades estimated using BI and ML above. The alignment was partitioned by codon and clock models and trees were linked in the three partitions. The site model and associated substitution model for each of the partitions was estimated using bModelTest (Bouckaert and Drummond, 2017). Other models such as strict versus relaxed log normal (Drummond et al., 2006) molecular clocks as well as Yule (Yule, 1925) versus birth-death (Gernhard, 2008) speciation models were compared using the steppingstone application in BEAST to generate marginal likelihood scores. Those models with the highest marginal likelihood estimate were used as priors in the final analysis. Prior calibrations included one fossil, the oldest *Spilogale* fossil (Hibbard, 1941a, b) and one molecular date for New World mephitids, (Eizirik et al., 2010) following Ferguson et al. (2017). Two separate runs of 50 million iterations each were sampled every 1,000 iterations. A burn-in of 10% was performed on each run and trees were combined using LogCombiner in BEAST. Effective sample sizes of posterior parameters were evaluated using Tracer v. 1.7 (Rambaut et al., 2018).

#### 2.5.2. Cytochrome *b* haplotype diversity analyses

We examined the extent of the distributional ranges of the putative species of *Spilogale* by examining all available sequences on GenBank for the cytochrome *b* gene, for which the most data are currently available (Ferguson et al., 2017; Shaffer et al., 2018) and for which genetic divergence has been shown to be useful for mammalian species-level comparisons (Baker and Bradley, 2006; Bradley and Baker, 2001). We visualized the cytochrome *b* haplotypes using a Median Joining Network (Bandelt et al., 1999) in the program POPART (Leigh and Bryant, 2015). Additionally, we estimated genetic distances between clades with the Kimura 2-parameter model (Kimura, 1980) using MEGA v. 6.06 (Tamura et al., 2013).

#### 2.5.3 Spatial diffusion modelling using continuous phylogeography

We used the unique mitochondrial haplotypes and corresponding geographic coordinates to generate an .xml file in BEAUTI v. 1.10.4 to perform the BEAST continuous phylogeography approach. The simple constant population size coalescent Gaussian Markov random field prior (Bayesian Skyride) was used as the demographic prior (Minin et al., 2008). Rate of diffusion was allowed to vary across branches by applying a Cauchy Relaxed Random Walk (RRW) model (Lemey et al., 2010). MCMC chains were run for 200 million generations, sampling every 10,000 generations. TREE ANNOTATOR v. 1.10.4 (Drummond et al., 2012) was used to summarize the posterior sample of trees including a 10% burn-in and to create a maximum clade credibility tree (MCC). TRACER v. 1.7 (Rambaut et al., 2018) was used to evaluate the adequacy of the 10% burn-in and to summarize the posterior sampling of each parameter. SPreaD3 v. 0.9.7.1rc (Bielejec et al., 2016) was used to visualize lineage diversification through time and space using the continuous diffusion model.

## 3. Results

Throughout the remainder of the manuscript, we use taxonomic names for the monophyletic groups following the taxonomic recommendations outlined in the Discussion section *4.5*.

### 3.1. Mitochondrial divergence and geographic structure

We recovered mitogenomes for 54 individual *Spilogale* specimens (Supplementary Table S1). The final mitogenome assemblies were similar in size, structure, and gene arrangement to other mammalian genomes. Part of the control region was removed from the final assemblies due to a repetitive region that made assembly difficult and therefore the final concatenated alignment was 16,345 nucleotides in length. We uncovered concordant evidence for geographically structured phylogenetic groups in both the concatenated ML analysis and with the partitioned dataset using BI (Fig. 1). We found support for eight monophyletic mitochondrial lineages that correspond to seven *Spilogale* species, including a highly divergent lineage, *S. pygmaea*, restricted to the Central Pacific Coast of Mexico. The relationships among the other taxa have been recovered in previous mitochondrial studies of *Spilogale* (Ferguson et al., 2017; Shaffer et al., 2018); however, in this study we recovered novel sequences of *S. angustifrons* and *S. yucatanensis* from historical museum specimens. Both ML and BI recover three major clades: *S. pygmaea*, an eastern clade, and a western clade. Within the eastern clade there is support for *S. putorius*, *S. interrupta*, and *S. yucatanensis*. The western clade contains support for *S. gracilis*, *S. leucoparia*, and *S. angustifrons*. We also recovered a unique mitochondrial clade we refer to as the Sonora Clade that is sister to *S. leucoparia* and *S. angustifrons* (referred to as the Arizona clade by Ferguson et al., 2017).

### 3.2. Multilocus nuclear phylogenies

Trinity assemblies produced an average of 21,732 contigs per sample (min = 6,449; max = 148,490). The grand mean for the assemblies was 489 nucleotides in length (min = 282; max = 672). We recovered 4,466 UCE loci in the incomplete matrix (N = 41 taxa; min 2119 and max = 3772). On average, the modern samples captured 3,296 loci, while the historical samples averaged 2,287 loci. We examined topologies with various levels of missing data: the 50% matrix contained 4,054 UCE loci, the 60% (i.e., 40% missing taxa per UCE locus) contained 3,896 loci, and the 70% matrix contained 3,511 loci. Our concatenated ML analysis estimated with the 60% matrix resolved most of the major clades; however, the relationship of *S. angustifrons* to *S. gracilis* and *S. leucoparia* was weakly supported (Supplementary Fig. 2).

The species tree analyses, using the three levels of missing data, estimated similar topologies with similar quartet values (Fig. 2, Supplementary Fig. 3). The species tree was concordant with the concatenated ML analysis indicating the support for the monophyletic relationship of the *S. angustifrons*, *S. gracilis*, and *S. leucoparia*; however, the relationship among the three taxa is not fully supported by all quartets (e.g., 88% of quartets resolved the sister relationship between *S. leucoparia* and *S. gracilis*). All other major clades were fully or highly supported in the species tree analysis.

**Figure 3.**
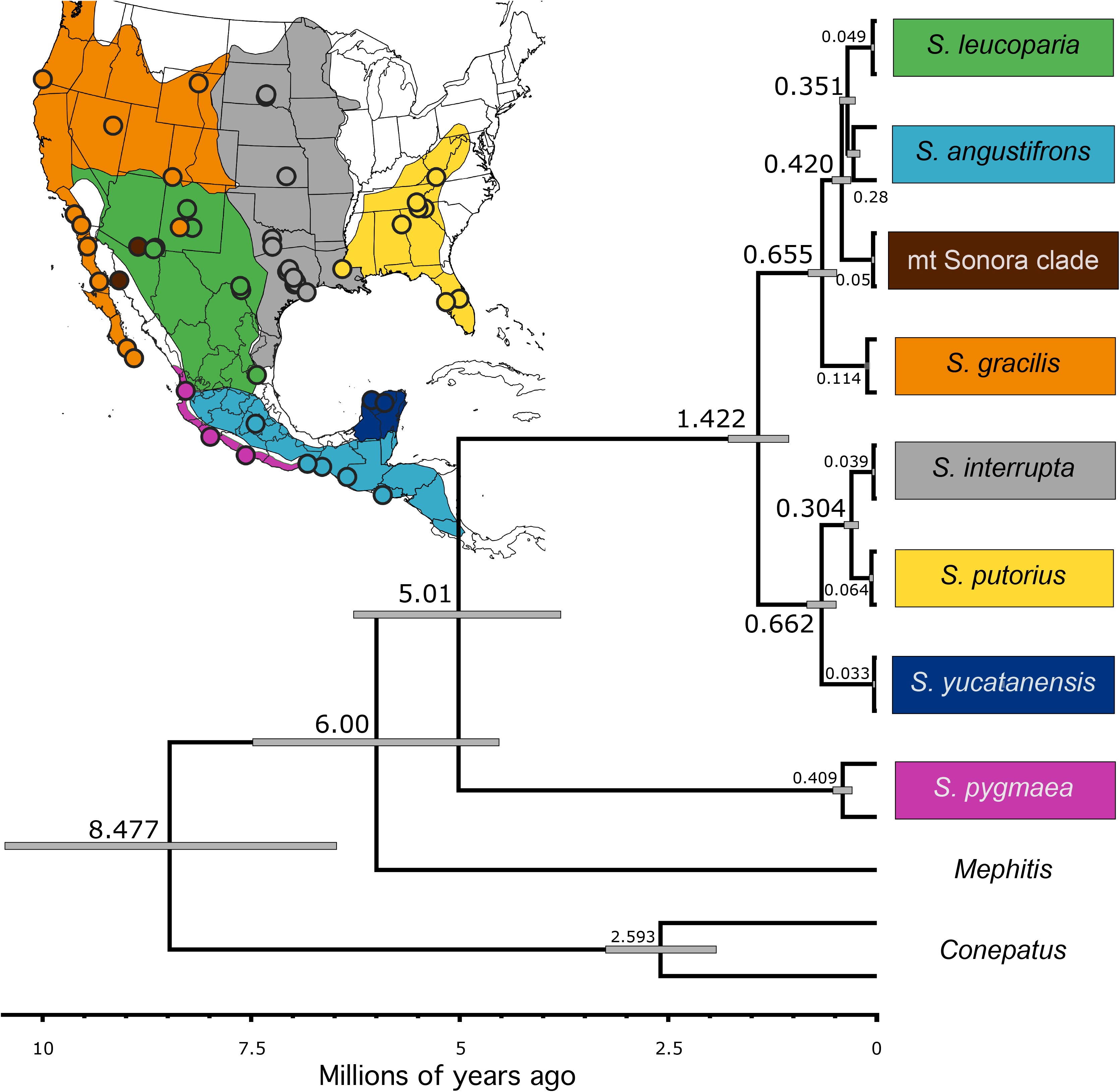
Divergence times estimated from 13 mitochondrial protein coding genes. Calibration priors included one fossil (Hibbard 1941, a and b) and one molecular date (Eizirik et al., 2010). Map inset includes sampling localities for mitogenomes (dots) and colored ranges estimated from cytochrome-*b* haplotype analysis.

Both the concatenated and species tree analyses failed to recover the distinct Sonora Clade, instead members from that clade were nested within *S. leucoparia*. We observed a further example of introgression with individual TK182721 which had *S. gracilis* mitochondrial DNA and *S. leucoparia* nuclear DNA.

### 3.3. Mitochondrial DNA analyses

#### 3.3.1. Divergence Time Estimation indicates an Early Pliocene origin of *Spilogale*

Our divergence time analysis that included the Yule speciation with a relaxed molecular clock had the highest marginal likelihood score compared to other models (Fig. 3). Our divergence times estimated using 13 mitochondrial protein coding genes and using one fossil calibration (Hibbard 1941, a and b) and one molecular calibration (Eizirik et al., 2010) estimated the origin for *Spilogale* occurred around 5 million years ago (95% highest posterior density [HPD] = 6.27 – 3.79 million years ago). The time to most recent common ancestor (TMRCA) for the eastern clades (*S. interrupta*, *S. putorius*, and *S. yucatanensis*) and western clades (*S. angustifrons*, *S. gracilis*, and *S. leucoparia*) occurred at roughly the same time: 0.66 million years ago (95% HPD = 0.84 – 0.49 million years ago and 95% HPD = 0.82 – 0.48 million years ago, respectively). The mitochondrial Sonora Clade diverged around 0.42 million years ago (95% HPD = 0.54 – 0.31 million years ago) followed by a nearly simultaneous final divergence between *S. angustifrons* + *S. leucoparia* and *S. putorius* + *S. interrupta* at 0.35 million years ago (95% HPD = 0.45 – 0.26 million years ago) and 0.30 million years ago (95% HPD = 0.39 – 0.22 million years ago), respectively.

#### 3.3.2. Haplotype diversity and mitochondrial cytochrome *b* divergence

We examined a subset of mitochondrial cytochrome *b* sequences from 203 individuals representing 139 unique localities to describe the haplotype diversity and geographic ranges of the taxa. These included 181 previously published sequences of the cytochrome *b* gene (Ferguson et al., 2017; Shaffer et al., 2018) and 22 novel cytochrome *b* sequences extracted from 18 whole mitogenomes used in the phylogenetic analyses and four extracted from historic samples with whole mitogenomes that were too poor quality for use in the mitogenome phylogenetic analyses but nonetheless contained high quality cytochrome *b* sequences (Supplementary Table S1). We recovered eight major haplogroups and 73 haplotypes that are concordant with the mitogenome ML and BI phylogenies (Supplementary Fig. 1). We recovered novel cytochrome *b* sequences for three Central American individuals (Costa Rica, El Salvador, and Nicaragua) that are 1.5% divergent from other *S. angustifrons* at the cytochrome *b* gene, but fall within the *S. angustifrons* group. Additionally, we found that the mt Sonora Clade is closely related to *S. angustifrons*.

#### 3.3.3. Spatial diffusion analysis and biogeographic history

We summarized the results of the spatial diffusion analysis at three timescales (Fig. 4). By 3.8 million years ago, the 80% HPDs of the ancestral locations contained two major regions: the southern United States and central Mexico + southern Mexico. By 1.5 million years ago, most of the geographic spread within *Spilogale* had occurred, including dispersal to the Yucatán. By 0.3 million years ago, MCC branches extend into the Baja Peninsula and the Great Plains.

**Figure 4.**
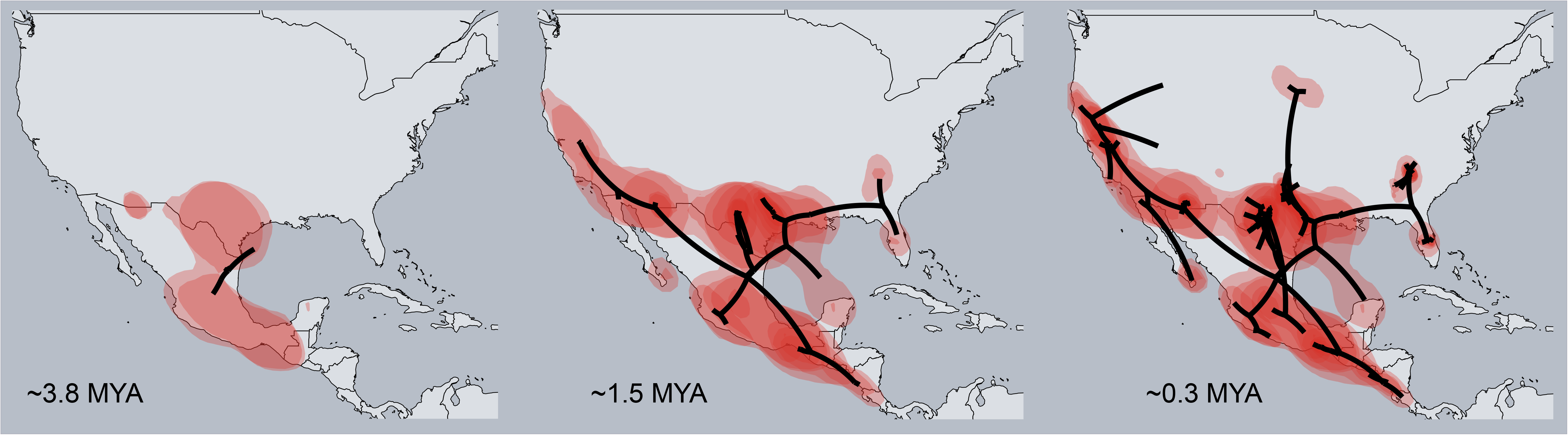
Estimates of spatial diffusion at three time periods: approximately 3.8, 1.5, and 0.3 million years ago. Red polygons indicate uncertainty (80% highest probability density) surrounding the geographic locations of the internal nodes of the tree and lines indicate maximum clade credibility (MCC) tree branches.

## 4. Discussion

### 4.1. Genetic diversity and structure

Our results demonstrate a significant degree of genetic differentiation within the genus *Spilogale* (Fig. 1; Fig. 2), indicating the presence of seven distinct evolutionary significant units, each reflecting distinct species (Fig. 3). Given the broad geographic range encompassing a number of distinct environmental conditions and complex geological features it is not surprising to find such genetically distinct populations for this group. However, the majority of this diversity appears to have arisen recently as a result of Pleistocene climate change associated with longer interglacial cycles. Generation of such diversity over a relatively short evolutionary timespan contradicts patterns seen in other, broadly distributed mephitids such as the striped skunk (Barton and Wisely, 2012) and North American hog-nosed skunk (Ferguson, 2014), but reflects similar patterns observed in other small carnivores with similar distributions (Harding and Dragoo, 2012; Nigenda-Morales et al., 2019).

Although the most recent mammal taxonomic compendiums recognize four species of spotted skunks (*S. angustifrons*, *S. gracilis*, *S. putorius*, and *S. pygmaea*; Wozencraft, 2005; Dragoo, 2009; Burgin et al., 2018) our phylogenetic analyses support the presence of seven species. This includes the splitting of *S. gracilis sensu stricto* into two distinct species, the Rocky Mountain spotted skunk *S. gracilis* confined to the western United States and Baja Peninsula which includes the currently recognized subspecies *S. g. gracilis*, *S. g. amphialus*, *S. g. latifrons*, *S. g. lucasana*, *S. g. martirensis*, and *S. phenax* and the desert spotted skunk *S. leucoparia* in the southern US and northern Mexico, which includes the entire distribution of the currently recognized subspecies *S. g. leucoparia*. The eastern spotted skunk *S. putorius sensu stricto* is also split into two distinct species, the prairie spotted skunk *S. interrupta* found in the central U.S. east of the Rocky Mountains and west of the Mississippi River and the Alleghany spotted skunk *S. putorius* which inhabits Appalachia and the southeastern US east of the Mississippi River, encompassing the range of two currently recognized subspecies *S. p. putorius* and *S. p. ambarvalis*. The southern spotted skunk, *S. angustifrons sensu stricto* also represents two species, the southern spotted skunk *S. angustifrons* found south of the Trans-Volcanic Mexican Belt into Central America but excluding the Yucatán Peninsula and encompassing four of the five recognized subspecies (*S. a. angustifrons*, *S. a. celeris*, *S. a. elata,* and *S. a. tropicalis*), and the endemic Yucatán spotted skunk *S. yucatanensis* which is elevated from the previously recognized subspecies *S. a. yucatanensis*. The pygmy spotted skunk *S. pygmaea* consists of a single species, rounding out the seven extant species of *Spilogale*.

The genetic characteristics of the pygmy spotted skunk *S. pygmaea* indicate that this species has long been isolated from other members of the genus (approximately 5 mya; Table 2). As such, *S. pygmaea* represents the most divergent lineage (Fig. 3), with an average of 15% divergence in the mitochondrial cytochrome *b* gene from other species. This level of divergence is more representative of generic differences among the mephitids rather than species-level differences (Ferguson et al., 2017). This degree of genetic differentiation is also reflected in the morphology of this diminutive species, which has morphological features more allied to the extinct Pleistocene *S. rexroadi* than to extant sister species (Dalquest, 1971). Such degree of morphological differentiation and genetic divergence warrants further investigation into the taxonomic assignment of the pygmy spotted skunk to the genus *Spilogale*.

**Table 2.** Divergence date estimates for time to most recent common ancestor for spotted skunk lineages (*Spilogale*) based on analysis conducted in BEAST v. 2.6.1 (Bouckaert et al., 2019) using a concatenated alignment of the 13 protein coding genes in the mitochondrial genome with significant biogeographic events and associated references indicated.

| MRCA | Ma | 95% HPD | Biogeographic Significant Events | References |
| --- | --- | --- | --- | --- |
| *S. pygmaea* + Other *Spilogale* | 5.01 | 6.27 – 3.79 | Sierra Madres and Trans-Mexican Volcanic Belt orogenesis- 5-35 Ma | (Ferrari et al., 1999) |
| “*S. gracilis*” + “*S. putorius*” | 1.42 | 1.78 – 1.06 | 6<sup>th</sup> NA glaciation<br>Cordilleran Ice Sheet advanced to outer limit of glaciation 1.5 Ma<br>7<sup>th</sup> NA glaciation | (Barendregt and Duk-Rodkin, 2011; Rutter et al., 2012; Westgate et al., 2001) |
| *S. yucatanensis* + *S. putorius*/*S. interrupta* | 0.66 | 0.84 – 0.49 | Shift from 41,000-100,000 periodicity cycle took place ~ 750 kya; 100 ky rhythm dominant by 600 kya | (Barnosky, 2005; Vrba, 1995) |
| *S. gracilis* + *S. leucoparia*/*S. angustifrons*/Sonora Clade | 0.66 | 0.82 – 0.48 | Increase in amplitude and periodicity of glacial-interglacial cycles ~800 kya ago | (Barnosky, 2005) and references therein |
| Sonora Clade + *S. angustifrons*/*S. leucoparia* | 0.42 | 0.54 – 0.31 | Mid-Brunhes event (MBE), roughly corresponds to the transition between stage 12 and stage 11 (Termination V) ~ 430 kya ago | (Augustin et al., 2004; Berger and Wefer, 2003) |
| *S. angustifrons* + *S. leucoparia* | 0.35 | 0.45 – 0.26 | Yarmouth Interglacial period (380-300 kya to 200 kya, pre-Illinoian stage) | (Voss, 1939) |
| *S. putorius* + *S. interrupta* | 0.30 | 0.39 – 0.22 | End of Reid Glaciation (0.13-0.28 Ma)<br>Start of Holstein Interglacial period (0.30-0.424 Ma) | (Barendregt and Duk-Rodkin, 2011; Rutter et al., 2012) |

The genetic divergence between western *S. gracilis* and eastern *S. putorius* (average of 7.24% divergence for cytochrome *b*), is supported by a longer period of isolation between the two species (Fig 3.; Supplementary Table S2) corresponding with the maximum glacial extent for the Cordilleran Ice Sheet (Westgate et al., 2001). The presence of delayed implantation in western populations of spotted skunks (Mead, 1968b) and absence in eastern populations (Mead, 1968a), indicating distinct species status based on the biological species concept (Mayr, 1942), accurately reflects the evolutionary distinctiveness of these two species. Interestingly, however, the two distinct eastern and western lineages of *Spilogale* are both composed of three distinct species each (Fig. 1; Fig 2; Table 2), with the western lineage including the currently recognized *S. angustifrons* and two distinct lineages corresponding to the western United States (*S. gracilis*) and southwestern US and north-central Mexico (*S. leucoparia*), respectively (Fig. 1; Fig. 2). Average between genetic divergences among the three western clades is 3.35% for the cytochrome *b* gene, with the lowest divergence values seen between *S. angustifrons* and *S. leucoparia* (2.42%) and *S. angustifrons* and the Sonora Clade (2.28%;) (Supplementary Table S2). The eastern lineage includes the endemic population of spotted skunks found in the Yucatán Peninsula (*S. yucatanensis*) and populations restricted to the central United States west of the Mississippi River (*S. interrupta*) and the eastern United States east of the Mississippi River through Appalachia and Florida (*S. putorius*). The genetic subdivision between subspecies of *S. putorius sensu stricto* was also recovered using microsatellite and mitochondrial markers (Shaffer et al., 2018). The three species within the eastern lineage shared an average cytochrome *b* divergence of 3.86% with the greatest difference observed between *S. yucatanensis* and *S. putorius* (4.87%). In the context of the larger dataset of 200 *Spilogale* samples, and solely focusing on the cytochrome *b* divergence results (>7%), a suggestion of just two species might be acceptable (Baker and Bradley, 2006), with the six lineages that make up these two divergent eastern and western clades representing values more in line with subspecies designation ~3.5% (Avise and Walker, 1999; Nigenda-Morales et al., 2019). However, this conclusion would be misguided given the results of our phylogenetic analysis based on both the UCE and mitogenome datasets. The greater number of informative characters from throughout the nuclear and mitochondrial genome yielded by these two markers provide the high statistical support needed for recognizing these six lineages as distinct species (Fig. 2; *4.5. Taxonomic recommendations* in Discussion).

Our molecular results are inconsistently congruent with historically proposed subspecies boundaries across the four currently recognized species of *Spilogale* (Dragoo, 2009), and as discussed in the preceding paragraph, are more reflective of unrecognized species-level diversity than misguided subspecies designations. The three recognized subspecies of *S. pygmaea*, *S. p. australis*, *S. p. intermedia*, and *S. p. pygmaea*, do seem to correspond to distinct clades in our phylogeny (Fig. 1; Fig. 2), although the limited sample size (n = 3) precludes definitive conclusions. The five recognized subspecies of *S. angustifrons* (*S. a. angustifrons*, *S. a. celeris*, *S. a. elata*, *S. a. tropicalis*, *S. a. yucatanensis*) appear to represent two distinct species, one corresponding to *S. a. yucatanensis* and the other by the four remaining subspecies. Interestingly, *S. a. yucatanensis* forms part of the phylogenetic clade including eastern populations of *S. putorius* instead of aligning with *S. angustifrons* and the western populations of *S. gracilis* (Fig. 1; Fig. 2). Phylogenetic results for the seven recognized subspecies of *S. gracilis* are similar to those observed in *S. angustifrons* with two of the currently recognized subspecies (*S. g. gracilis* and *S. g. leucoparia*) corresponding to distinct species, and molecular support for recognizing three of the remaining subspecies as valid: *S. g. amphialus*. *S. g. lucasana*, and *S. g. martirensis*. Although not represented in our UCE or mitogenome phylogenies, previous phylogenetic analyses based on mtDNA and microsatellite data indicate that the island spotted skunk *S. g. amphialus* warrants subspecific recognition (Ferguson et al., 2017; Floyd et al., 2011). Interestingly, the genetically distinct mitochondrial lineage referred to as the Sonora Clade does not correspond to any historically recognized metric of diversity, although it clearly is reflective of an historically isolated population dating back to the Pleistocene (Ferguson et al., 2017). For the three subspecies of *S. putorius* (*S. p. ambarvalis*, *S. p. interrupta*, and *S. p. putorius*), two correspond to distinct species (*S. p. interrupta* and *S. p. putorius*). The third, *S. p. ambarvalis* from Florida, nested within the clade including *S. p. putorius* from Appalachia and other southeast coastal states of Alabama, Georgia, and Louisiana (east of the Mississippi River), and as such was not supported as a distinct subspecies.

The small size, and likely limited dispersal capability, and high mortality rate of spotted skunks (Hillman et al., 2014; Jones et al., 2013; Lesmeister et al., 2010) may help explain the strong population structure observed in both the mitochondrial and nuclear data, especially when compared to patterns observed in other skunk species (Barton and Wisely, 2012; Ferguson, 2014) and other small carnivores such as the red fox, *Vulpes vulpes* (Aubry et al., 2009; Volkmann et al., 2015) and ermine, *Mustela erminea* (Dawson et al., 2014). Patterns observed in *Spilogale* appear more consistent with other codistributed small mammals such as bats (Weyandt and Van Den Bussche, 2007) and rodents (Mantooth et al., 2013; Nava-García et al., 2016), although a similar pattern of divergence and genetic diversity was found in the long-tailed weasel, *Mustela frenata* (Harding and Dragoo, 2012). Similar to the major geographic subdivisions observed in *Spilogale*, five major mitochondrial clades were recovered for *M. frenata*, including a western North America, eastern North America, north-central Mexico, southern Mexico-Central America, and South America clade, with similar biogeographic barriers implicated in limiting gene flow between these clades (i.e., the Mississippi River, Trans-Mexican Volcanic Belt, and North American deserts (Harding and Dragoo, 2012).

### 4.2. Phylogeographic and divergence patterns

A majority of the observed genetic subdivisions appear to more strongly correspond with Quaternary climate fluctuations than pre-Pleistocene barriers, although a number of known barriers do coincide with boundaries of proposed ranges of the phylogenetic clades. The divergence between the pygmy spotted skunk and all other species of *Spilogale* around 5 million years ago appears to correspond with the completion of the orogenesis of the Trans-Mexican Volcanic Belt and Sierra Madres during the late Miocene and early Pliocene (Ferrari et al., 1999; Ferrari et al., 2005). Other significant physiographic barriers to gene flow include the Mississippi River, a pattern shared with other small carnivores such as the raccoon *Procyon lotor* (Cullingham et al., 2008) and striped skunk (Barton and Wisely, 2012), the Vizcaino Seaway of the Baja Peninsula (Riddle and Hafner, 2006), and the Mojave Desert and Boues Embayment (Riddle and Hafner, 2006). The Sierra Madre Occidental and Oriental appear to restrict *S. leucoparia* to the Mexican Plateau region, limiting gene flow between populations in the Sonoran Desert and Gulf Coast of Mexico.

Of particular interest is the lack of phylogroups associated with other major biogeographic barriers known to impact mammals, including other skunk species. For instance, the Sierra Nevada Mountains appear to have had little to no impact in generating distinct phylogroups for *Spilogale,* in comparison to patterns observed in striped skunks (Barton and Wisely, 2012) and deer mice *Peromyscus* (Yang and Kenagy, 2009). And although our limited sampling across Central America restricts our ability to draw inferences for populations inhabiting this area, the cytochrome *b* data which includes samples from throughout the region, indicates a lack of phylogroups associated with major biogeographic barriers observed for other, co-distributed small mammals (Gutiérrez-García and Vázquez-Domínguez, 2012), with the exception of the Yucatán Peninsula (Vázquez-Dominguez and Arita, 2010). In addition, the Isthmus of Tehuantepec, which has acted as a major biogeographic barrier for a number of small mammals (Ordóñez-Garza et al., 2010) appears to have had a limited isolating role for *Spilogale*.

The high levels of genomic divergence between the seven major phylogroups, or species (See *4.5. Taxonomic recommendations*), appear to have mostly resulted from glacial-interglacial cycles during the Pleistocene (Fig. 3, Table 2). The initial divergence into the two major clades, an eastern and western clade, corresponds to major glaciation periods in North America at the time, including the maximum extent of Cordilleran Ice Sheet around 1.5 million years ago (Westgate et al., 2001). Subsequent divisions of these two major lineages, both into two distinct phylogroups per lineage, occur at nearly the same time, around 0.66 million years ago (Fig. 3). This estimate reflects a significant change in the glacial cycles that began around 0.74 million years ago and continued through 0.43 million years ago, with a transition to a higher proportion of each cycle spent in the warm mode (Augustin et al., 2004), and alteration of the periodicity cycle from a rhythm dominated by interglacial cycles of 0.04 million years ago to one of 0.10 million years ago (Barnosky, 2005). This change in glacial cycles meant longer periods of higher sea levels, warmer temperatures, and increased CO_2_ production during interglacial periods, a pattern that held for nearly every 0.10 million years during the interglacial cycle that began 0.80 million years ago (Hansen et al., 2013).

The Sonoran Clade which maintains a distinct mitochondrial lineage but a shared nuclear genome with *S. leucoparia* diverged approximately 0.42 million years ago, corresponding to the mid-Brunhes event (Berger and Wefer, 2003), a period of climate change that reflects changes in temperatures and greenhouse gases similar to Earth’s present interglacial period (Augustin et al., 2004; Berger and Wefer, 2003). Although not tested in the present paper, paleo-ecological niche models indicated that this mitochondrially distinct phylogroup was isolated to one or two climate refugia in northern Sonora during the last interglacial (Ferguson et al., 2017).

The final subdivision of the broader eastern and western lineages, into two distinct species each, also occurred at nearly the same time (0.35 – 0.30 million years ago). Within the western lineage, the split between *S. angustifrons* and *S. leucoparia* appears to correspond with the Yarmouth interglacial period and that between *S. putorius* and *S. interrupta* of the eastern lineage with the start of the Holstein interglacial period. These patterns of coincident subdivision of the two major western and eastern lineages indicates a shared process of isolation and diversification during glacial maximum followed by range expansion during interglacial periods.

These climate-induced subdivisions of populations also have significant implications for our understanding of the evolution of delayed implantation in spotted skunks. Although strong evidence of the presence or absence of delayed implantation is restricted to populations of *S. gracilis* and *S. putorius sensu stricto* (Mead, 1968a; Mead, 1968b), the fact that these two species are sympatric in various parts of their range (e.g., the two species co-occur in the same small geographic area in Central Texas; Dowler et al., 2008), supports the notion that delayed implantation acts as an effective pre-zygotic isolating mechanism. Given the historical split between eastern and western lineages of *Spilogale* prior to subsequent diversification within each of these lineages, one might hypothesize that the common ancestor from the tropical south would lack delayed implantation, which is generally considered an adaptation to highly seasonal environments with extreme periods of low productivity (Ferguson et al., 2006; Ferguson et al., 1996; Mead, 1989; Thom et al., 2004). Then, following subsequent isolation of this historical population into two distinct lineages due to expanded glacial coverage, especially for the western US, western lineage populations might have evolved delayed implantation. If so, one might expect delayed implantation in the three species that compose the western lineage, *S. angustifrons*, *S. gracilis*, *and S. leucoparia*, but not in those of the eastern lineage, *S. interrupta, S. putorius, and S. yucatanensis*. Data from Mead (1968a, 1968b) documented the presence of delayed implantation in *S. gracilis* and its absence in *S. interrupta* and *S. putorius* from Florida (*S. p. ambarvalis sensu stricto*) with anecdotal accounts indicating its presence in *S. leucoparia* (Constantine, 1961) and absence in *S. angustifrons* from Central America (Creed and Biggers, 1964) and *S. yucatanensis* (Van Gelder, 1959). Further investigations on the reproductive biology of populations from central Mexico, Central America, the northern Great Plains, and Appalachia would shed light on the origin and evolution of delayed implantation in *Spilogale*.

### 4.3. Introgression and secondary contact

The number of studies reporting mito-nuclear discordance appears to be increasing, likely reflecting the simultaneous use of a suite of genetic markers across broad taxonomic and geographic sampling regimes (Riddle and Jezkova, 2019; Toews and Brelsford, 2012). For *Spilogale*, a pattern of mito-nuclear discordance, or more aptly, biogeographic discordance (Toews and Brelsford, 2012), is supported by the presence of eight mtDNA clades in comparison to seven clades based on nuclear DNA alone (Fig 1; Fig. 2). The biogeographically restricted Sonora Clade recovered using mtDNA only (Fig 3; see also Ferguson et al., 2017) appears to have been isolated since the Last Interglacial and bound by well-characterized pre-Pleistocene barriers, despite its much more recent origins (Ferguson et al., 2017). Despite this historic isolation, the Sonora Clade is either currently in secondary contact with, or at some point in the past was in contact with, the *S. leucoparia* clade in southwestern Arizona (Fig. 1), perhaps across the Cochise filter barrier (Ferguson et al., 2017; Morafka, 1977). We uncovered two specific instances of mitochondrial introgression for the Sonora Clade with *S. leucoparia* both of which were from specimens captured in Arizona’s Pima County (ASK5307 and HD73). Interestingly, these two occurrences of introgression were found in a region of the densest sampling for our study, with 24 samples sequenced for mtDNA in southern Arizona (Fig. 1). The second most densely sampled area of west-central Texas (n = 18 sequences of mtDNA, Fig. 1) did not recover evidence of introgression, although this region represents the contact zone between the much older lineages of western and eastern *Spilogale*, thought to be isolated by the presence and absence of delayed implantation, respectively (Mead, 1968a; Mead, 1968b). The geographic region inhabited by the Sonora Clade roughly corresponds with a northwest Sonoran clade of banded geckos (*Coleonyx variegatus sonoriensis*), which experienced introgression with a distinct species inhabiting southwest Arizona during the Late Quaternary (Leavitt et al., 2020). Our results support the call from Leavitt et al., (2020) for increased sampling across the Sonoran Desert to further elucidate patterns of isolation and secondary contact among vertebrates inhabiting this region.

A second instance of mito-nuclear discordance was observed in southern New Mexico, involving the western *S. gracilis* clade and east-central *S. leucoparia* clade. This sample from Lincoln County, New Mexico (TK182721) possessed a mitogenome that clearly aligned with *S. gracilis* but a nuclear genome that belonged to the *S. leucoparia* clade (Fig. 1). In contrast, a second sample, collected less than 1 km away (TK22414), possessed both a mitochondrial and nuclear genome assigned to *S. leucoparia*. The individual displaying patterns of introgression is the only sample of *S. gracilis* found east of the Rio Grande River, a known biogeographic barrier for many southwestern vertebrates (Riddle and Hafner, 2006). Although our limited sampling across northern Arizona and New Mexico limits our inferences regarding the extent of this secondary contact, our results suggest that gene flow has been present between *S. gracilis* and *S. leucoparia*. Increased sampling across Arizona and New Mexico, and other regions where secondary contact has been documented in other co-distributed taxa (e.g., across the Mexican highlands of the TMVB; Zarza et al. 2018), coupled with whole genome sequencing would certainly help elucidate historical versus contemporary patterns of gene flow among the distinct lineages of *Spilogale* (Colella et al., 2018).

### 4.4. Biogeographic history/Ancestral range reconstruction

New World skunks are thought to have arisen from a single immigration event from Eurasia via Beringia, reaching the west coast of North America in the late Clarendonian (Wang et al., 2005). The oldest skunk taxon, *Martinogale faulli*, dated to the Late Miocene’s Dove Spring Formation (~ 10 million years ago), is thought to have given rise to a transitional form in western Texas in the early Hemphillian, spreading into the Great Plains in the late Hemphillian (Wang et al., 2005). The earliest fossil records of *Spilogale* appear in the upper Pliocene (3.5 – 3 million years ago) from late Blancan sites in Kansas and Texas (Wang et al., 2014; Wang et al., 2005). The species represented at both sites, *S. rexroadi* was a “diminutive skunk”, similar in size to the living *S. pygmaea* and smaller races of *Spilogale*, e.g., *S. a. yucatanensis,* (Dalquest, 1971). Although incomplete, the skunk fossil record indicates that southern North America, including Mexico and Central America, acted as the center for diversification of *Spilogale* (Wang et al., 2014). Prior to the discovery of additional skunk fossil material, Van Gelder (1959) proposed that the distribution of extant *Spilogale* species was the result of post-glacial recolonization of western and eastern North America from ancestral populations found in southern United States and Mexico. Both Van Gelder (1959) and Mead (1989) also pointed to the increased diversity of *Spilogale* in Central America and Mexico as evidence for a southern North American origin of the genus.

Results from our spatial diffusion analysis indicate a northeastern Mexico and southern US (Texas) origin of *Spilogale* (Fig. 4) supporting these previous hypotheses regarding a southern North American origin for the genus (Mead, 1989; Van Gelder, 1959; Wang et al., 2005). Although isolation of distinct isoclines based on our samples appear as early as 4 million years ago, the expansion out of these isolated core ranges appears to reach a peak during the interglacial periods around 0.3 million years ago, in conjunction with the final diversification into the seven distinct clades. Of particular interest is the early identification of an isolated population in the Yucatán Peninsula, with a likely dispersal path from northeastern Mexico/southern Texas along Mexico’s Gulf Coast (Fig. 4). Although a lack of samples from Mexico’s Gulf Coast limits our ability to draw firm conclusions, adjacent populations to *S. yucatanensis* in northeastern Mexico and Central America belong to the western linages *S. angustifrons* and *S. leucoparia* instead of its closest relative, *S. putorius*, which inhabits the southeastern United States including southern Florida. This may suggest the existence of a tropical biogeographic corridor connecting the southeastern United States and northeastern Mexico to southern Mexico and Central America during Plio-Pleistocene glacial intervals (Morgan and Emslie, 2010). We found that *S. yucatanensis* is more closely related to fauna in northern Mexico/southeastern United States. This is in contrast to what other studies have found where other small mammals of the northern Yucatán which are closely related to fauna in the southern Yucatán Peninsula (Guevara et al., 2014; Gutiérrez-García and Vázquez-Domínguez, 2012; Nigenda-Morales et al., 2019; Vázquez-Dominguez and Arita, 2010). This could be due to similar habitats shared between the two regions (Morrone et al., 2002) or historical connections that existed during periods of lower sea levels and extensive corridors of tropical habitats (Morgan and Emslie, 2010). Determining the range limits of *S. yucatanensis*, which adds another endemic species of vertebrate to this biogeographic region already characterized by high levels of endemism (Vázquez-Dominguez and Arita, 2010), would require samples from the Petén region of the peninsula in countries like Belize and Guatemala.

### 4.5. Taxonomic recommendations

Although a full taxonomic revision is beyond the scope of this manuscript, given the conservation implications for currently recognized *Spilogale* taxa (USFWS, 2012; Gompper 2017), the overlap or intergradation seen in morphology across these taxonomic boundaries (Van Gelder, 1959), and the clear patterns for species level recognition seen in our molecular datasets, assigning names to the identified clades is important to help recognize and protect the cryptic diversity observed in spotted skunks. Although a complete taxonomic revision, including multiple lines of evidence (e.g., comparative morphology, karyotypes, and ecology) is warranted and necessary, the need for immediate recognition of the existing diversity in *Spilogale* seems prudent. Thus, based on historical descriptions of species including their purported geographic distributions and the principle of priority (Ride et al., 1999), we propose the following taxonomic arrangement to describe the molecularly identified clades of *Spilogale*.

#### *Spilogale leucoparia* Merriam, 1890-Desert spotted skunk

*Spilogale leucoparia* Merriam, 1890, N. Amer. Fauna, 4:11.—Type Locality. “Mason, Mason County, Texas”. Holotype. USNM 186452, skull, skin adult male, collected by I. B. Henry on 02 December 1885.

<u>Original Description</u>: “White markings larger than any other known species, the white on back equaling or even exceeding the black in area’ all the stripes are broader than in the other species; the middle pair of dorsal stripes are continuous posteriorly with the anterior transverse stripe, which in turn are broadly confluent with the extern lateral stripes. The lumbar spots are generally confluent with the posterior transverse stripes. The tail spots are sometimes confluent posteriorly, forming a narrow band across the base of the tail. There is no white on the thighs, and only rarely a few white hairs on the upper surface of the foot.”

<u>Original Range</u>: None provided

<u>Interpreted Range</u>: The desert spotted skunk, *S. leucoparia* appears to be limited in its distribution to the Chihuahuan and Sonoran Deserts, including the Madrean Sky Islands (Coronel-Arellano et al., 2018), but it also is found throughout the Mexican Plateau (Escalante et al., 2003). The proposed range appears to be bound to the north by temperate grasslands associated with the southern Rocky Mountains, the south by the Trans-Mexican Volcanic Belt (Ferrari et al., 1999), the west by the Colorado River and Mojave Desert, and the east by the Great Plains. The presence of individuals belonging to this clade in high elevation pine-oak forests could partially explain the individual sample from Gomez Farias, which is within the Tamaulipan biotic province, and support the presence of this lineage in either the Sierra Madre Occidental or Oriental. Further sampling of individuals along the Gulf Coast of Mexico, northeastern Mexico, and southern Texas is needed to further define the range limits of *Spilogale* species in this region.

#### *Spilogale gracilis* Merriam, 1890-Rocky Mountain spotted skunk

*Spilogale gracilis* Merriam, 1890, N. Amer. Fauna, 3:83.—Type Locality. “Grand Canon of the Colorado (altitude 3,500 feet), [Coconino County, 1067 m] Arizona, N of San Francisco Mountain.” Holotype. USNM 17986, skin, skull, adult male, collected by C. H. Merriam and V. O. Bailey on 12 September 1889.

<u>Original Description</u>: Longer and more slender than the eastern *S. putorius,* with a much longer tail. Frontal white patch much longer than broad, and rounded off both above and below; dorsal and lateral markings essentially as in *S. putorius*. Terminal part of the tail white, the white occupying a little more than a third of the upper surface and two-thirds of the under surface.

<u>Original Range</u>: “The Little Striped skunks are characteristic members of the Sonoran fauna, and do not occur at higher altitudes than this fauna or its offshoots attain. They are rarely found from far water, and most of the species prefer rocky situations, often making their homes in crevices in cliffs.”

<u>Interpreted Range</u>: The Rocky Mountain spotted skunk, *S. gracilis*, appears to inhabit much of the western United States and the entire Baja Peninsula of Mexico, including the Great Basin Desert, and north into the temperate rainforests of the Pacific Northwest. The proposed range appears bound to the north by grasslands of the Great Plains and the Canadian Rocky Mountains, the south by the Colorado Plateau, the west by the Pacific Ocean, and the east by the Great Plains. Further sampling of individuals along the zone of contact between *S. leucoparia* and *S. gracilis* in northern Arizona and New Mexico is needed to define the range limits of *Spilogale* species in this region.

#### Subspecific Taxonomy

##### *Spilogale gracilis amphialus* Dickey, 1929-Island spotted skunk

*Spilogale phenax amphialus* Dickey, 1929, Proc. Biol. Soc. Washington, 42:158.—Type Locality. “2½ miles north of ranch house near coast, Santa Rosa Island, Santa Barbara County, California”. Holotype. UCLA 13400, skin, skull, adult male, collected by H. H. Sheldon on 6 November 1927.

<u>Original Description</u>: “Externally the island animal has the short total length of *microrhina*, but differs strikingly from it in having the tail not long as in that form, but even shorter both actually and relatively than the short-tailed northern form *phenax*.”

<u>Original Range</u>: “Santa Rosa and Santa Cruz Islands, Santa Barbara County, California. Association, chiefly cactus patches, on Santa Rosa at least.”

<u>Interpreted Range</u>: The island spotted skunk is restricted to the Channel Islands of Santa Barbara County, California, including the islands of Santa Cruz and Santa Rosa.

##### *Spilogale gracilis gracilis* Merriam, 1890-Rocky Mountain spotted skunk

*Spilogale gracilis* Merriam, 1890, N. Amer. Fauna, 3:83.—Type Locality. “Grand Canon of the Colorado (altitude 3,500 feet), [Coconino County, 1067 m] Arizona, N of San Francisco Mountain.” Holotype. USNM 17986, skin, skull, adult male, collected by C. H. Merriam and V. O. Bailey on 12 September 1889.

<u>Original Description</u>: See species account above.

<u>Original Range</u>: See species account above.

<u>Interpreted Range</u>: The Rocky Mountain spotted skunk, *S. gracilis gracilis* appears to inhabit much of the western United States, excluding the entire Baja Peninsula of Mexico and the Channel Islands of California. The proposed range of this subspecies appears bound to the north by grasslands of the Great Plains and the Canadian Rocky Mountains, the south by the Colorado Plateau, the west by the Pacific Ocean, and the east by the Great Plains‥

##### *Spilogale gracilis lucasana* Merriam, 1890-Southern Baja spotted skunk

*Spilogale lucasana* Merriam, 1890, N. Amer. Fanua, 4:11.—Type Locality. “From Cape St. Lucas, Lower California”. Holotype. USNM 3970 skin, skull, adult male, collected by John Xantus.

<u>Original Description</u>: “Size large; tail long (with hairs apparently about as long as head and body); terminal pencil white; white markings large and broad.” “This is the only species known to me in which there is any regularity in the throat and chin markings.” “…[skulls] are much larger, broader posteriorly, flatter, and everywhere more massive than those of any other species examined.”

<u>Original Range</u>; None provide.

<u>Interpreted Range</u>: The range of the subspecies *S. g. lucasana* appears to be restricted to Mexico’s Baja California Sur, the northern boundary of which appears to correspond with the location of the historical barriers of the Vizcaino Seaway and Isthmus of La Paz (Riddle and Hafner, 2006). Further sampling in the northern portions of Baja California Sur is needed to determine the exact range limits of *S. g. lucasana* across the Peninsula.

##### *Spilogale gracilis martirensis* Elliot, 1903-Northern Baja spotted skunk

*Spilogale arizonae martirensis* Elliot, 1903, Field Columb. Mus., Publ. 74 Zool. Ser., 3:170.— Type Locality. “Vallecitos, San Pedro Martir mountains, Lower California, 9,000 feet elevation.” Holotype. FMNH 10572, skin, skull, male, collected by E. H. Heller on 23 September 1902.

<u>Original Description</u>: “Similar to *S. arizonae* in markings, but the white stripes from occiput and cheeks narrower and shorter, those from cheeks reaching only to just beyond shoulders’ broken stripe from fore leg across lower back, broader’ tail and hind foot short. Skull is shorter, narrower, and lighter, with narrower rostrum.”

<u>Original Range</u>: None provided.

<u>Interpreted Range</u>: The range of the subspecies *S. g. martirensis* appears to be restricted to Mexico’s Baja California, the southern boundary of which appears to correspond with the location of the historical barriers of the Vizcaino Seaway (Riddle and Hafner, 2006).

##### *Spilogale angustifrons* Howell, 1902-Southern spotted skunk

*Spilogale angustifrons* Howell, 1902, Proc. Biol. Soc. Wash. 15:242.—Type Locality. “Tlalpam, Valley of Mexico”. Holotype. USNM 50825, skin, skull, adult male, collected by E. W. Nelson and E. A. Goldman on 15 December 1892.

<u>Original Description</u>: “Size small; coloration as in *S. ambigua,* but usually without the white bands on the thighs. Skull slender, and without prominent ridges.” “This form belongs with the group of narrow-skulled species inhabiting the eastern United States, in which group *ambigua* also belongs.”

<u>Original Range</u>: “The present form occupies the southern portion of the Mexican table-land, from Guanajuato to Chiapas.”

<u>Interpreted Range</u>: The southern spotted skunk *S. angustifrons* appears to inhabit the southern half of Mexico (excluding the Yucatán Peninsula) south to Costa Rica. The proposed range appears bound to the north by Trans-Mexican Volcanic Belt, the south by the North Panama Fracture Belt (Gutiérrez-García and Vázquez-Domínguez, 2012) the west by the Mexico’s

Pacific dry forest, and the east by Veracruz moist forests. Additional samples from the coastal Mexican states of Veracruz, Tabasco, and Campeche are needed to fully understand the eastern limits of this species.

##### *Spilogale yucatanensis* Burt, 1938-Yucatán spotted skunk

*Spilogale angustifrons yucatanensis* Burt, 1938, Occ. Pap. Mus. Zool. Univ. Mich., 384:1.— Type Locality. “Chichen Itzá, Yucatán, Mexico”. Holotype. UMMZ 75780, skin, skull, adult female, collected by M. B. Trautman on 2 March 1936.

<u>Original Description</u>: Described by Burt (1938) as “a dwarf race of *angustifrons*” differing from *S. a. elata* and other races of *S. angustifrons* “in much smaller size, smaller teeth, and narrow tooth row.” Although small, it is described as differing from *S. pygmaea “…*in color pattern, having black instead of white feet, and in more inflated mastoids and more triangular as well as larger skull.”

<u>Original Range</u>: “Probably ranges over most of the Yucatán Peninsula.”

<u>Interpreted Range</u>: The Yucatán spotted skunk *S. yucatanensis* appears to be endemic to the Yucatán Moist and Dry forest habitats of the northern half of the Yucatán Peninsula, although our limited sampling from this region restricts our ability to define the species range boundaries. Sampling the Petén and Veracruz Moist forest habitats in Campeche, Belize, and northern Guatemala are required to define the limits of this species.

##### *Spilogale interrupta* (Rafinesque, 1820)-Plains spotted skunk

*Mephitis interrupta* Rafinesque, 1820, Ann. Nat., 1:3.—Type Locality. “Louisiana”. Holotype. None designated.

<u>Original Description</u>: “Brown, with two short parallel white streaks on the head, and eight on the back, the four anterior ones equal and parallel, and the four posterior ones rectangular, angles in opposite directions.”

<u>Original Range</u>: “…inhabiting Louisiana.” The exact location and source material for Rafinesque’s description of *Mephitis interrupta* is very lacking, a theme that seems to permeate many of his mammal species descriptions (Woodman, 2015, 2016). Lichtenstein took Rafinesque’s reference to this species “…inhabiting Louisiana” to mean the author must have been referring to the Louisiana Territory, and as such restricted the range of this species to the “Upper Missouri River Valley” (Lichtenstein, 1838: 281). Hall and Kelson (1959) further supported this restriction, recognizing the type locality as “Upper Missouri [River?]”.

<u>Interpreted Range</u>: The plains spotted skunk *S. interrupta* appears to be mostly restricted to the Great Plains of the United States although its range limits extend into the forested Ouachita and Ozark Mountains and south into the coastal prairies and marshes of Texas and parts of Louisiana (west of the Mississippi River). The proposed range of this species is bound to the north by southern Canadian forests, the south by the Tamaulipan biotic province (see interpreted range of *S. leucoparia*), the west by the Rocky Mountains, and the east by the Mississippi River, which has functioned as historical barrier to gene flow between *S. interrupta* and *S. putorius*.

##### *Spilogale putorius* (Linnaeus, 1758)-Alleghanian spotted skunk

[*Viverra] Putorius* Linnaeus, 1758, Syst. Nat., 10th ed., 1:44.— Type Locality. “America septentrionali” restricted by Thomas (1911a) and Howell (1901) to “South Carolina.” [USA]. Holotype. Based on Catesby (1774) “pol-cat” T. 62 illustration from Vol II, part 9 of Catesby’s Natural History of Carolina, Florida & the Bahama Islands (1739-1771)

<u>Original Description</u>: “V. fulsca lineis quatuor albidis dorsalibus parallelis. Cauda longitudine corporis, villosa. Irritata ut antecedens halitum explodit, quo nihil foetidius.”

<u>Original Range</u>: “Mississippi, Alabama, western Georgia, western South Carolina, and northward along the Alleghenies to northern Virginia; western limits of range unknown” according to (Howell, 1906).

<u>Interpreted Range</u>: The Alleghanian spotted skunk *S. putorius* inhabits the Alleghanian biotic subregion (Escalante et al., 2013), including most of the Appalachian Mountains and the southeastern coastline of the United States, including the entire state of Florida. The proposed range of this species is bound to the north by the northern piedmont and Allegheny plateau, the south by the Gulf of Mexico, the west by the Mississippi River, and the east by the piedmont and Atlantic Coastal Plain.

##### *Spilogale pygmaea* Thomas, 1897-Pygmy spotted skunk

*Spilogale pygmaea* Thomas, 1897, Proc. Zool. Soc. Lond., 1897: 898.— Type Locality. “Rosario, Sinaloa, W. Mexico”. Holotype. BMNH 98.3.2.24., skin, skull, adult sex unknown, collected by P. O. Wilson on 2 April 1897.

<u>Original Description</u>: “Size very small, barely half that of any known species. Pattern of coloration differing considerably from that found in the other members of the genus,…”. White of forehead united to the white ear-patches so as to form a band across the face from ear to ear, but in the centre of the face the white did not project forward beyond the level of the eye.” “Upper surfaces of both hands and feed white, in continuation in front with the white lateral stripe, and behind with a white line running up on the hams;

<u>Original Range</u>: Western Mexico.

<u>Interpreted Range</u>: The pygmy spotted skunk *S. pygmaea* is endemic to the tropical dry forests of Mexico’s Pacific lowlands, from Sinaloa south to Oaxaca (Ceballos et al., 2014). The recorded range appears to be restricted in the north by the Sonoran Desert, the south by the Isthmus of Tehuantepec, the west by the Pacific Ocean, and the east by the broadleaf and pine forests along the Sierra Madre del Sur and Trans-Mexican Volcanic Belt.

## 5. Conclusions

Our study reveals that the genetic diversification of *Spilogale* originated in the southern United States/northeastern Mexico during the Pliocene but that species level diversity within the genus was mostly driven by glacial-interglacial cycles coincident with a shift toward longer periods of interglacial cycles around 0.70 – 0.80 million years ago. Phylogenetic analyses coupled with geographic range limits indicate the presence of seven extant species of *Spilogale*: *S. angustifrons, S. gracilis, S. interrupta, S. leucoparia, S. putorius, S. pygmaea,* and *S. yucatanensis*. These results support a complex evolutionary history with examples of secondary contact between historically isolated populations across the Desert Southwest. These findings have direct implications for our understanding of delayed implantation and for informed conservation management within *Spilogale*. Additional sampling is needed along the Gulf Coast of Mexico, across Central America, and in northern Arizona and New Mexico to comprehensively understand the extent of secondary contact and exact distributional limits for several of the recognized species.

## Acknowledgements

This manuscript fulfilled partial requirements for the PhD degree to A.W.F. under the supervision of R. E. Strauss with special thanks to committee members R. J. Baker, R. D. Bradley, and A. Townsend Peterson. A suite of modern collectors, curators, and museum staff deserve special recognition for facilitating access to tissue samples for genetic assays including T. Lee of Abilene Christian University Natural History Collection, L.K. Ammerman, M. Revelez, D. Krejsa, J.C. Perkins, and A. Shaffer of the Angelo State Natural History Collection, D. Spaulding of Anniston Museum of Natural History, B. Sasse of Arkansas Game and Fish Comission, R. Birkhead of Auburn University, J. Light of the Biodiversity Research and Teaching Collections at Texas A&M University, S. Miller of the Bob and Betsy Campbell Museum of Natural History, Carolina Museum of Science, F. X. González-Cózatl and E. Arellano Arenas of the Centro de Investigación en Biodiversidad y Conservación Universidad Autónoma del Estado de Morelos, S. T. Alvarez Castañeda of the Centro de Investigaciones Biológicas del Noroeste, C. Lorenzo of the Colección Mastozoológica de El Colegio de la Frontera Sur, R. Eng, and D. Jachowski of Clemson University, J. Demboski and J. Stephenson of the Denver Museum of Nature and Science, L. Heaney and B. Patterson of the Field Museum of Natural History, T. Hannon and the Florida Fish and Wildlife Conservation Commission, V. Mathis of the Florida Museum of Natural History, N. Castleberry of the Georgia Museum of Natural History, C. Lopéz González and D. Garcia of the Instituto Politecnico Nacional Colección Cientifica de Fauna Silvestre, M. Gabriel of the Integral Ecology Research Center and the University of California Davis, L. L. Panguia of the Museo de Zoología “Alfonso L. Herrera” de la Facultad de Ciencias de la Universidad Nacional Autónoma de México, J. Cook and J. Dunnum of the Museum of Southwestern Biology at the University of New Mexico, I. E. Engilis and A. Hitch of the Museum of Wildlife and Fish Biology at University of California Davis, H. Garner and K. McDonald of the Natural Sciences Research Laboratory at Texas Tech University, J. Frey of New Mexico State University, G. Albers and C. Olfenbuttel of North Carolina Wildlife Resources Commission, S. Tremor of the San Diego Natural History Museum, D. Lunde and S. Peurach of the Smithsonian Institution’s National Museum of Natural History, C.J. Schmidt of the Sternberg Museum at Fort Hays State University, M. Rheude, M. Culver, and T. Edwards of the University of Arizona, D. Van Vuren of the University of California Davis, M. Eifler and R. M. Timm of the University of Kansas Biodiversity Institute and Natural History Museum, J. Krupa of the University of Kentucky, C. W. Thompson of the University of Michigan Museum of Zoology, J. W. Dragoo of the University of New Mexico, E. Rickart of the Utah Museum of Natural History, M. Fies of Virginia Department of Game & Inland Fisheries, N. Moncrief of the Virginia Museum of Natural History, A. Edelman and T. Sprayberry of West Georgia University. The following individuals also provided tissue samples C. Cornelison, D. Leismeister, J. Herbst, P. Eyheralde, D. Reding, V. Evelsizer, and J. Strickland. This research was conducted using the Smithsonian Institution High Performance Cluster (SI/HPC): https://doi.org/10.25572/SIHPC. This project and the preparation of this publication was funded in part by the State Wildlife Grants Program (Grant #AT-T0F16AF01278) of the U.S. Fish and Wildlife Service through an agreement with the Arkansas Game and Fish Commission. Additional funding came from research grants and scholarships to A.W.F. from the American Museum of Natural History, American Society of Mammalogists, and the Southwestern Association of Naturalists.

---

**Supplementary Figure 1.**
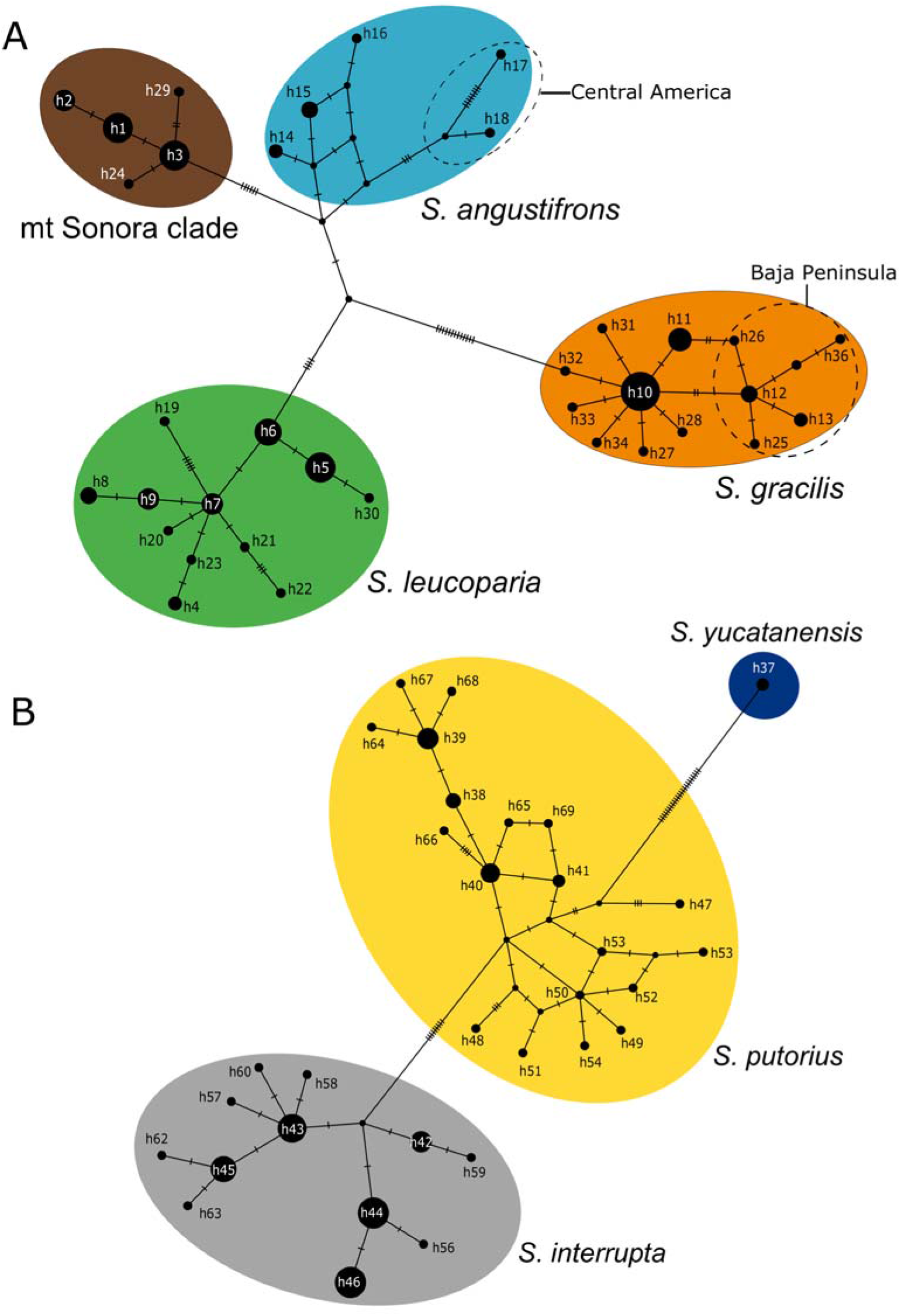
Median-joining networks generated from the mitochondrial cytochrome-*b* gene for (A) “eastern group” and (B) “western group”. Subgroups of interest include individuals from the Baja Peninsula and the divergent individuals from Central America shown with dashed line.

**Supplementary Figure 2.**
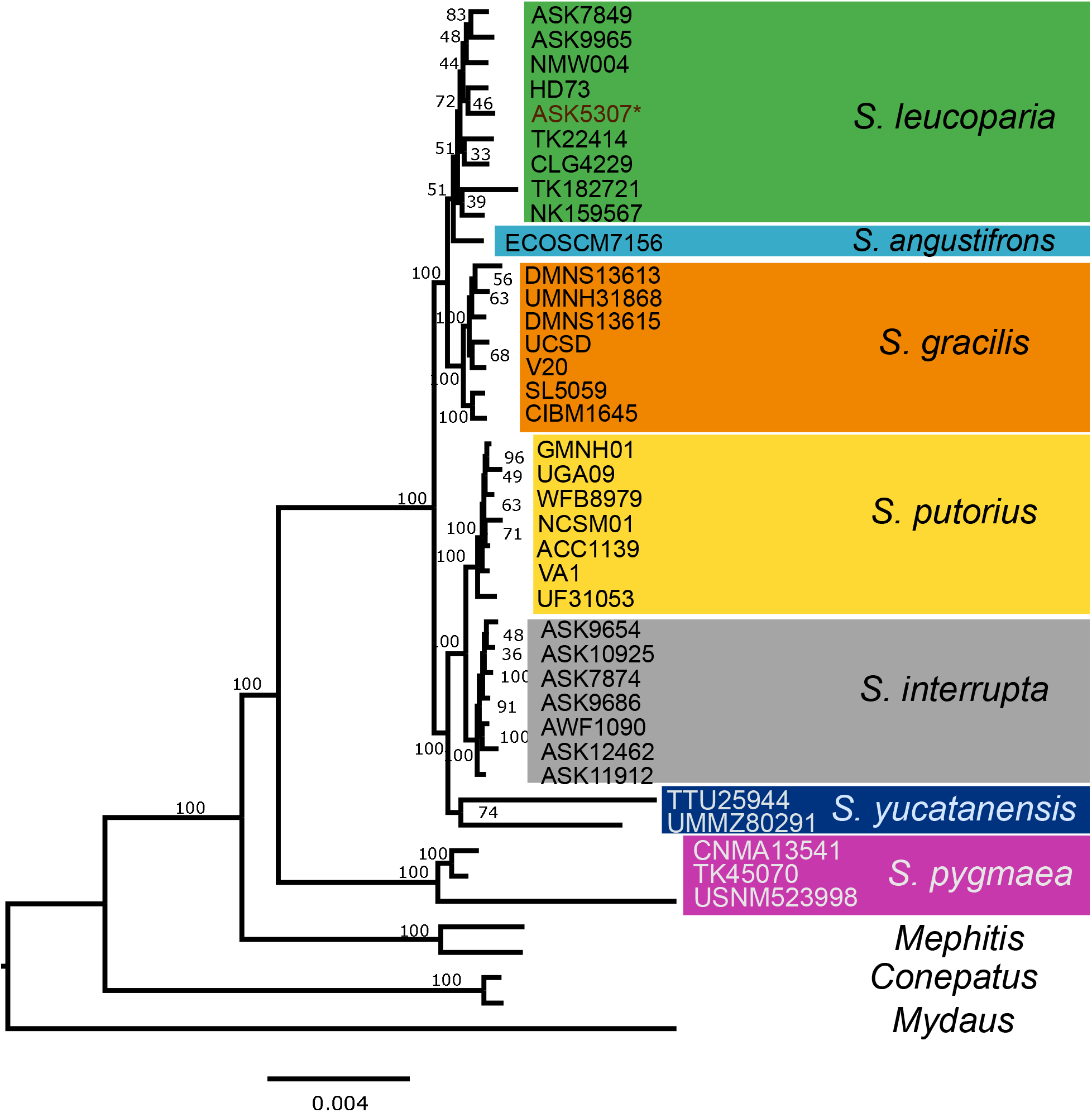
Maximum-likelihood analysis of *Spilogale* species based on analysis of 3,896 UCE loci (60% matrix). Bootstrap values estimated from 500 replicates.

**Supplementary Figure 3.**
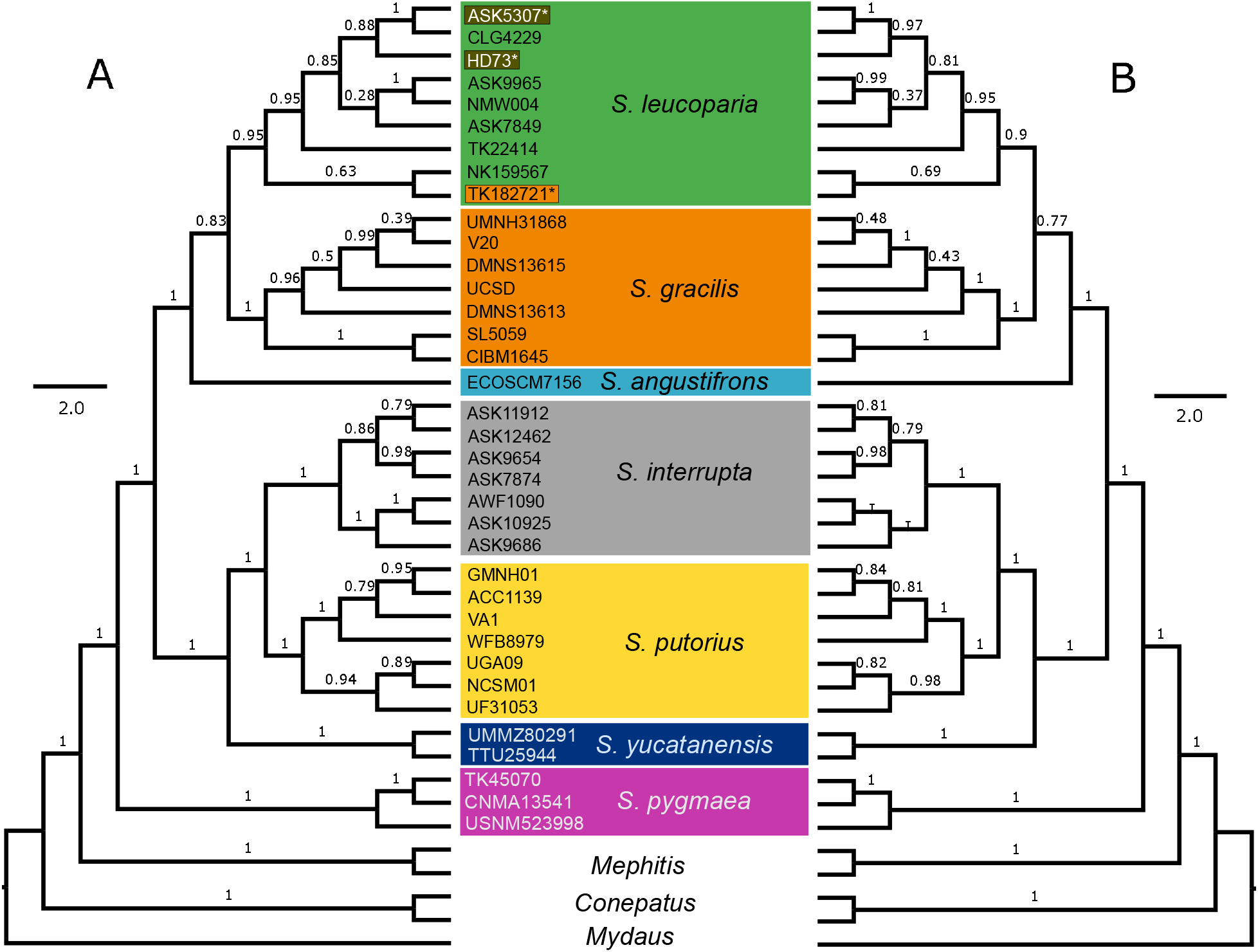
Cladogram of *Spilogale* species based on analysis of A) 4,054 UCE loci UCE loci (50% matrix) and B) 3,511 loci (70% matrix). Quartet values from species tree are displayed along the branches.

